# Contrasting Impacts of the Anthropogenic Environment on the Diet of Long-tailed Macaques (*Macaca fascicularis*) in Southeast Asia

**DOI:** 10.1101/2025.07.26.666990

**Authors:** Benjamin Gombash, Amanda Acevedo, Chissa Rivaldi, Hope Hollocher

**Author notes:** Correspondence: Benjamin Gombash 520 Westgate Ave, Apartment 302, University City, MO, 63130 1-419-345-9949.

## Abstract

Human expansion will only increase access to anthropogenic food resources for wildlife species. Access to these resources can have positive and negative impacts on wildlife species. It is generally assumed that access to anthropogenic resources will decrease diet diversity, although recent work questions this. Our research uses metabarcoding to assess the diet detected in fecal samples from long-tailed macaques (*Macaca fascicularis*) on the islands of Singapore and Bali, Indonesia. We gathered information about multiple parameters to characterize the distinctly different anthropogenic environments on the two islands. Then, we used a leave one out cross validation model selection procedure and linear regressions to determine which environmental parameters impact various diet metrics for the macaques in these two different anthropogenic contexts. Macaques in both island contexts increased their usage of crop diet items with greater anthropogenic access, but with different resulting patterns. We found that as food provisioning of the macaques increases in Bali, multiple measures of diet richness decrease, as generally expected, yet diet stability within a group also decreases, indicating that although diet richness decreases with provisioning, individuals do not experience similar levels of access to these resources. In contrast to Bali, there were no significant differences in diet stability across groups in Singapore with increases in anthropogenic resources. These differing impacts on diet, seen in a single species, highlight the need to consider more closely the details of the spatial and temporal distribution of anthropogenic resource access when making predictions about how a species’ diet will be impacted.

## Introduction

As human settlement expands in many geographical contexts, anthropogenic food resources are becoming a more common element of the diet of many species of wildlife. Anthropogenic resources can take multiple forms that are relevant to wildlife, such as intentional provisioning, crop-foraging, and refuse scavenging (Hill, 2018; McKinney, 2015).

Anthropogenic food items are generally high calorie, spatially clumped, temporally predictable, and abundant (Griffin et al., 2017; Sha & Hanya, 2013), allowing organisms that exploit them to benefit. However, there are some potential detriments associated with anthropogenic resources, such as reduced dietary taxonomic diversity and specific nutrient deficiencies (Murray et al., 2015; Griffin et al., 2017). Additionally, anthropogenic resource access can alter behaviors (Chauhan and Pirta, 2010; Sussman et al., 2011; Jaman and Huffman, 2013), activity budgets (O’Leary and Fa, 1993; Boug et al., 1994; Ilham et al., 2018), and parasite exposure and transmission patterns (Weyher et al., 2006; Lane et al., 2011; Becker et al., 2017; Becker et al., 2018). Specific dietary traits can impact how a wildlife species responds to anthropogenic resource access. Dietary generalists, which can utilize a wide range of diet items, often take advantage of anthropogenic landscapes more successfully than dietary specialists, which can only exploit a narrow range of diet items (Lowry et al., 2013; Fehlmann et al., 2020). Despite these general trends, the details of a species and the anthropogenic resources being utilized can have unexpected impacts on its diet. For example, urban coyotes actually exhibited more diverse diets than rural coyotes in contrast to expectations, which was likely due to coyotes exploiting novel food sources in some cases and broadening their diet to acquire protein that is lacking in anthropogenic foods in other cases (Murray et al., 2015). A meta-analysis that focused on the diets of vertebrate predators found that fish, birds, and reptiles showed the expected decrease in dietary richness in urban areas, which include greater access to anthropogenic resources, as compared to other habitats, although the effect was not significant. Most mammals and amphibians in this study saw an increase in dietary richness in urban environments, which likely reflects more generalist diets, although this effect was also non-significant. Therefore, the overall trend for all taxa was that there was, in fact, no significant relationship between predator dietary richness and urban environments (Gamez et al., 2022). However, this non-significance could be due to context-specific effects of different factors that are cancelled out across such broad datasets. Investigating how the diet of a single generalist wildlife species changes across diverse habitats with varying levels of anthropogenic resource access could help us identify the most important parameters influencing diet responses within anthropogenic contexts. These parameter-specific associations between specific kinds of anthropogenic resources and diet response may be more predictable across systems than looking for one general pattern between broadly described anthropogenic habitats and dietary richness.

Our research investigates interactions between the anthropogenic landscape and the diet of long-tailed macaques on the islands of Bali, Indonesia, and Singapore. Long-tailed macaques are widely distributed and are found in a diverse array of habitat types across their range, including multiple anthropogenic habitats, such as temples, rural settlements, urban areas, agricultural areas, and roadsides (Ong and Richardson, 2008; Gumert, 2011). Long-tailed macaques live in multi-male, multi-female groups and have a social system marked by intense unilateral aggression without frequent reconciliation, which can make mates and resources easy to monopolize. Like other macaque species, females are the philopatric sex, which results in matrilineal hierarchies where entire matrilines outrank each other. Males disperse, and may challenge a dominant male upon joining a group to gain access to mating opportunities, or they may form coalitions to overthrow dominant males in other contexts (Thierry, 2011; Fuentes et al., 2011). They are primarily frugivorous, but also eat insects, grass, leaves, flowers, mollusks, and fungi (Yeager, 1996; Wheatley, 1980; Lucas and Corlett, 1998; Brotcorne, 2014; Gumert, 2011). The long-tailed macaque is one of the most arboreal macaques; however, they are known increase their terrestrial behavior in anthropogenic habitats (Thierry, 2011; Brotcorne, 2014). Observations suggest that macaques tend to focus their feeding efforts on a limited subset of their diet (Wheatley, 1980; Yeager, 1996; Fuentes et al., 2011). This could result in a diet that is has two subsets, one of regularly consumed diet items and one of less regularly consumed items.

On the island of Bali, long-tailed macaques are known to inhabit a range of habitats, including agricultural and urban areas (Fuentes et al., 2011; Lane et al., 2011; Southern, 2002). These macaques have religious significance for the Hindus of Bali and have a sacred status while on temple grounds and possibly in other contexts (Louden et al., 2006; Lane de Graaf et al., 2013). Different types of food offerings are made at temples, sometimes to fulfill religious obligations, and sometimes to provision macaques. The volume and frequency of these offerings vary across the island (Fuentes et al., 2011; Lane et al., 2011). Nature-based tourism, which includes visiting temples to see macaques, has become an important part of Bali’s tourism-based economy (Sutawa, 2012). Temples that are visited more frequently tend to have more provisioning which has allowed macaque populations to increase in number (Lane-de Graaf et al., 2014).

In contrast to Bali, Singapore is largely urban with a central catchment area that is partially occupied by a nature reserve (Sha & Hanya, 2013). Despite it being illegal, humans sometimes directly provision macaques by throwing food from cars (Sha and Hanya, 2013; Fuentes et al., 2008), although there is no direct provisioning by the government or park officials (Fuentes et al., 2008). Given the close proximity to densely populated urban areas, the government of Singapore has utilized culling to control macaque population sizes in the past (Riley et al., 2015). Previous work in Singapore suggests that macaques inhabiting more anthropogenically altered environments have higher diet diversity, which was attributed to lower food availability forcing the macaques to spread out and forage across a larger area, exposing them to a wider range of diet items (Sha & Hanya, 2013). It is possible that this more dispersed foraging of a more diverse diet could have resulted in lower diet similarity within the group of macaques, although this was not specifically investigated. Exploring how within group diet similarity responds to anthropogenic resource access may provide valuable insights into how a species responds to these resources.

In order to assess macaque diet broadly across these two contrasting habitats, we utilized a metabarcoding approach, which entails using high throughput sequencing to identify many taxa from a sample containing the DNA of many species (Soininen et al., 2014). These techniques have been used previously to investigate different aspects of animal diets, including diet plasticity related to landscape variation (Quemere et al., 2013), anthropogenic contributions to the diet (Forin-Wiart et al., 2018), and, more broadly, diet composition (Soininen et al, 2013; Deagle et al., 2018, Albainia et al., 2016; Barba et al., 2014; Mollot et al., 2014; Leray et al., 2015; Pompanon et al., 2012; Tournayre et al., 2020). These approaches have multiple advantages over more traditional methods when it comes to diet assessment, such as being less time intensive, allowing for a broader sampling of individuals, and potentially identifying diet items more specifically (Quemere et al., 2013). Metabarcoding techniques can provide higher resolution diet assessment when foraging for diet items that are difficult to observe with traditional methods, such as arthropod foraging by arboreal primates (Lyke et al., 2019), and can reveal a broader dietary range than was previously detected (McLennan et al., 2022). One disadvantage of metabarcoding approaches is that they cannot distinguish which part of an organism was consumed (e.g. fruit or leaves). Beyond measurements of diet richness, similarity metrics have been used to investigate diet overlap within and between groups of organisms (Mower and Smith, 1989; Storms et al., 2008), and can be used as a proxy for evaluating the stability of the diet across a group. Here we include this metric to determine how the distribution and/or access to diet items are impacted in different anthropogenic contexts. It is not known whether changes in diet richness (i.e. the number of items in the diet) necessarily result in changes in diet stability, or consistency, across individuals within a group, as this has not been investigated previously. A wider sampling of individuals, facilitated by our metabarcoding approach, can help to define core and secondary diets, which can demonstrate foraging preferences or differences in availability between diet items (Tournayre et al., 2020).

Our research here addresses multiple questions about how anthropogenic resources influence the diet of long-tailed macaques. First and foremost, we are interested in understanding whether anthropogenic resource access affects macaque diet richness. Although the assumed expected pattern is to see a decrease in diet richness with an increase in anthropogenic resource utilization (Gamez et al., 2022), we anticipate that we will recapitulate previous results in Singapore (Sha & Hanya, 2013) and see an increase in diet richness with increased anthropogenic resource access. We expect the opposite in Bali, where regular active provisioning (Lane et al., 2011) makes anthropogenic resources more predictable. Second, we investigate the role of crop plants, a group of diet items that have an anthropogenic origin, in the long-tailed macaque diet. We predict that macaques with more anthropogenic resource access will utilize more crop plants in their diet and rely more heavily on crop plants. Third, we assess whether anthropogenic resource access impacts diet similarity, or stability, among a group of macaques. We expect the diet to become more stable across a site in Bali, where anthropogenic resources are predictable, but less stable in Singapore, where anthropogenic resources are less predictable (Sha & Hanya, 2013). We will determine how the subset of diet items that are most prevalent in a group of macaques, their majority diet, respond to anthropogenic resource access. We anticipate that the majority diet of macaques in Singapore will decrease in richness due to the unpredictable nature of anthropogenic resources, which is in line with our predictions about increasing overall diet richness and decreasing stability. For macaques in Bali, we predict that the majority diet will increase in richness as the overall diet becomes less rich and more stable. The different traits associated with the anthropogenic resources in Bali and Singapore produce different predictions for how anthropogenic resources impact the macaques’ diets.

## Methods

### Ethical Note

All of this work was conducted with previously collected samples (Citations removed for anonymity). Sample collection was approved by the (Institution Name Removed for Anonymity) Institutional Animal Care and Use Committee (protocol IDs 07-001, 14-002, 14-05-1835). Samples collected from Bali were approved by the Indonesian Institute of Science (permit number 662.02/1090.DIII). Samples collected from Singapore had approval of the National Parks Board (permit #NP/RP11-029).

### Fecal Sample Collection and DNA Extraction

Fecal samples were collected from eight sites across Singapore (n = 63) from 2011 to 2013 (Supplemental Table 1; Klegarth, 2015) and twelve sites across Bali (n = 64) in the summer of 2007 (Lane et al., 2011; Figure 1; Table 1). In brief, fresh samples were collected from each site on a single day to avoid sampling an individual multiple times. Some defecation events were observed, but not all, and individual macaques were not identified. A part of each sample was kept on ice (Bali) or dry ice (Singapore) until they could be frozen at -85° for molecular analysis (Klegarth et al., 2017; Lane et al., 2011; Lane-deGraaf et al., 2014). DNA was extracted from each sample using the Qiagen QIAMP Stool Minikit (Qiagen, Hilden, Germany) following the manufacturer instructions (Qiagen, 2012; Wilcox and Hollocher, 2018).

**Figure 1:**
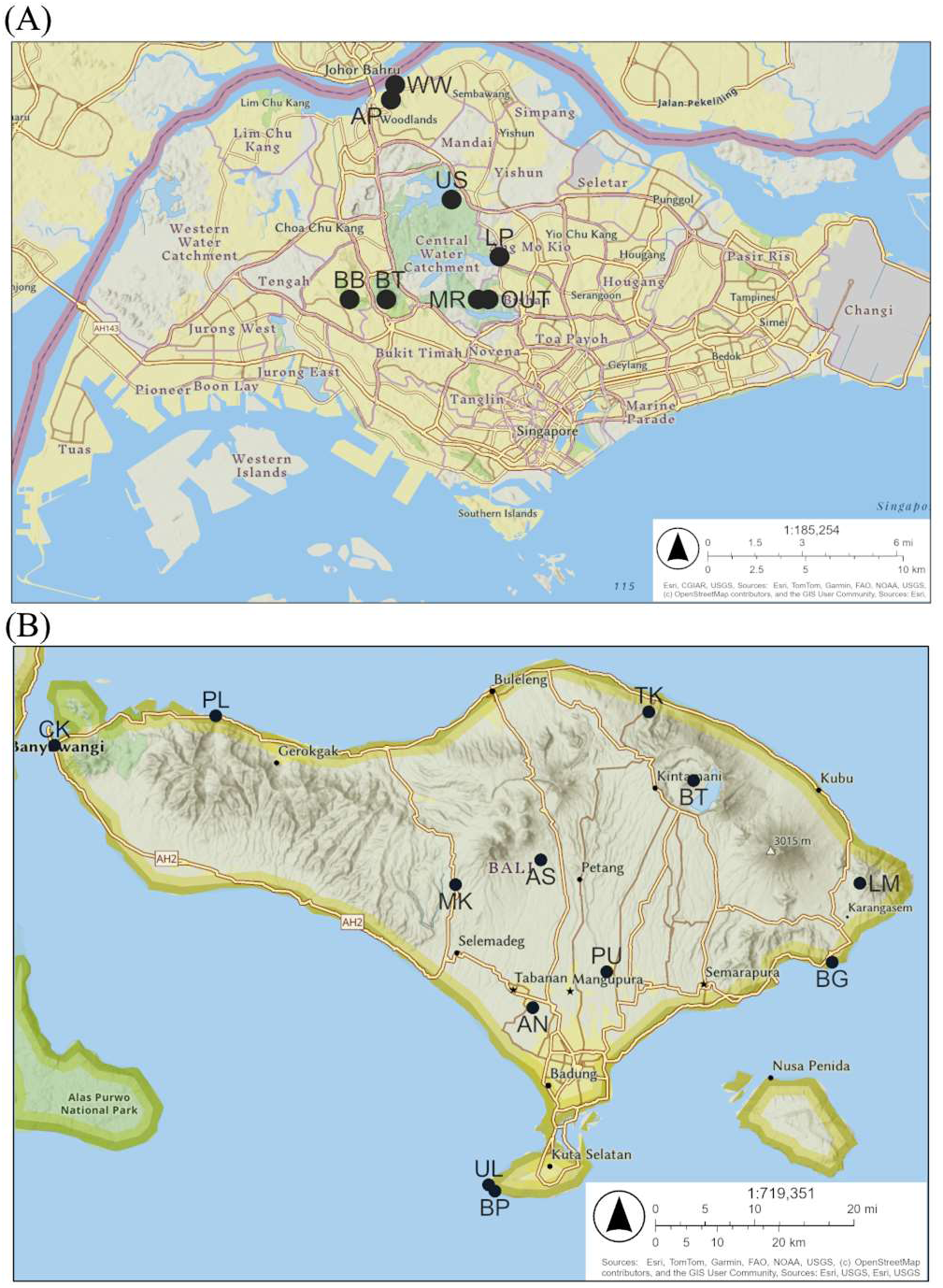
**(A)** A map of sites from Singapore where fecal samples and environmental descriptions were collected. **(B)** A map of sites from Bali where fecal samples and environmental descriptions were collected. Maps were generated in ArcMap Online utilizing the National Geographic basemap (Esri, 2025; National Geographic, 2025).

**Table 1.**
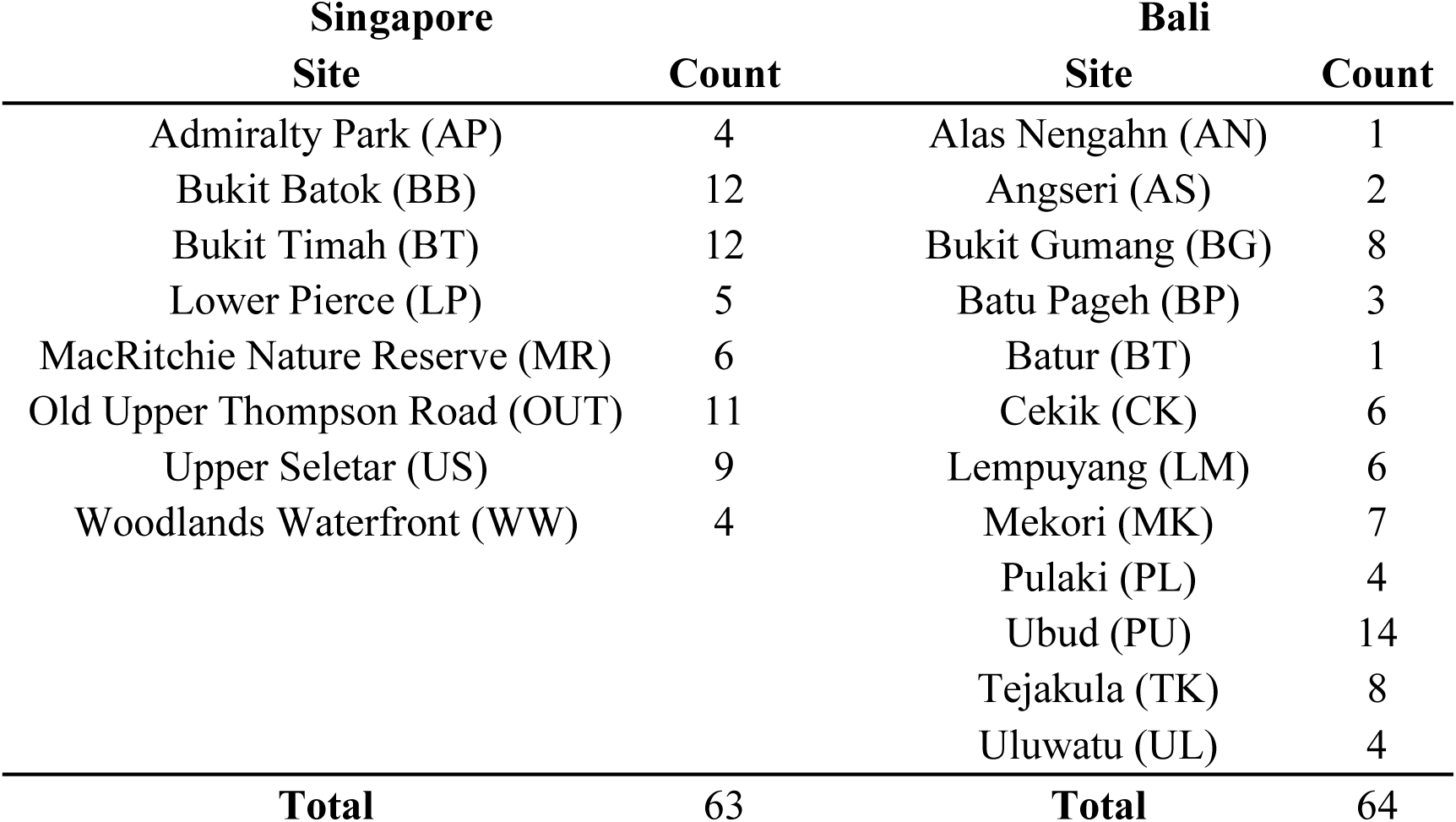
Sample counts from Singapore and Bali broken down by site.

### Target Amplification and High-Throughput Sequencing 18S ssu rRNA

Metabarcoding techniques have been used to investigate the diet, with the *trn*L being a popular gene amplicon to investigate the plant-based diet items (Kartzinel et al., 2015). For this research, the *18S ssu rRNA* gene amplicon was chosen because it can be used to capture a wider range of eukaryotic taxa, including plants and animals (Wilcox and Hollocher, 2018). The amplicon sequencing data used in this research were generated by two different sequencing runs, both of which used the procedure outlined in Wilcox and Hollocher (2018). A 90 to 180 bp sequence of the V9 hypervariable region of the *18S small-subunit ribosomal RNA* gene was amplified from genomic DNA using two forward primers (1380F and 1389F). These two primers share a reverse primer (1510R), which does not amplify macaque sequences, which eliminates the need for blocking primers. The two forward primers were used to capture a wider range of taxa than either primer could capture alone (Wilcox and Hollocher, 2018; Amaral-Zettler et al., 2009). The use of two forward primers that share a reverse primer prevents amplicon sequence variants (ASVs) from being created.

### Illumina PCR and sequencing and Bioinformatics Analysis

PCR products were purified, the libraries pooled, and afterwards the libraries were sequenced on an Illumina HiSeq 2500 (Illumina, San Diego, CA, USA) following the procedure outlined in Wilcox and Hollocher (2018). Data were analyzed following a modified USEARCH workflow (Wilcox and Hollocher, 2018; Rivaldi, 2022). One notable change is that the Silva 132 was used rather than Silva 123 (Quast et al., 2013). This process generated an *18S* operational taxonomic unit (OTU) table.

### OTU Dietary Genera Table Preparation

OTUs from the following taxa were considered potential diet items: the embryophyte clade, the phylum Mollusca, or the phylum Arthropoda. These taxa are considered *potential* diet items because we do not have observations to confirm consumption, but macaques are known to eat taxa from these groups (Brotcorne, 2014; Gumert, 2011; Lucas and Corlett, 1998; Wheatley, 1980; Yeager, 1996). OTUs with sufficient taxonomic information were amalgamated to the genus resolution. The data were filtered following a method outlined by Cirtwill and Hamback (2021) to remove potential false positive detections. We tested cut off values from one to twenty-five, where cells with fewer reads than the cut off value were set to zero. The number of links (i.e. non-zero cells in the table) kept and the percentage of links kept were calculated for each of the tested cut off values. These values were plotted, and these plots were used to select a cut off value of six reads to filter the potential dietary genera table. This means that any cell that had fewer than six reads was set to have zero reads (Supplemental Methods and Figure S1).

### Statistical Analyses Calculating Diet Metrics

Dietary genera richness (hereafter dietary richness) values were calculated for each sample by counting the total number of relevant dietary genera present in the sample. Overall dietary richness and crop genera richness (hereafter crop richness) were calculated. Genera were considered “crops” if a literature search recovered evidence that the genus was cultivated for human consumption as food. This means that genera such as *Allium* (onions and garlic), *Zea* (maize), and *Solanum* (potatoes and tomatoes) were considered crops whereas other cultivated genera such as *Nicotiana* (tobacco) and *Gossypium* (cotton) were not. We did not limit the search to crops that are known to be cultivated on Bali or Singapore given potentially imported foods would still have an anthropogenic origin. Similarly, we are not suggesting that the macaques gained access to these crop foods by raiding agricultural fields. For example, crop foods could be foraged from refuse bins or provisioned. Due to uneven sampling across sites, richness values were averaged across samples from each site.

Diet stability, or the consistency of the diet across samples at a site, was calculated using diet similarity following Ricotta and Pavoine (2015) (Equation 1). The variable *N* represents the total number of samples from a given site, *S* represents the total number of genera detected in samples from a given site, and *Ni* represents the number of occurrences within samples from a given site of dietary genus *i*. It is noteworthy that you cannot calculate similarity with a single sample, which means that we cannot calculate similarity for site BT on Bali.

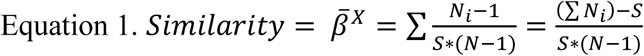

Using prevalence thresholds to determine whether a taxon is part of a core is common practice in metabarcoding studies, with thresholds ranging from 30% to 100%, although higher thresholds are more common (Neu et al., 2021). This technique is used in metabarcoding studies of the diet (Tournayre et al., 2020). We are using the term majority diet genera to describe any dietary genus that appeared in at least 50% samples from a given site. Linear regressions were performed to ensure that variable sample counts were not significantly related to the number of majority dietary genera for each site (Supplemental Figure S2). Employing a 50% threshold means that sites with two or fewer samples (AS, AN, and BT on Bali) are not altered by this process.

### Environmental Information

The anthropogenic environment of Bali was assessed previously by Southern (2002) and Lane et al. (2011) using a mixture of GIS data and personal observation (Supplemental Table 2). Information was collected about the landscape, paying special attention to forested area, urban area, rice agricultural area, and elevation. Other factors, such as tourism days, a measure of provisioning (offering weight in kg/day), and days with available water to drink (i.e. water days) were recorded as well. Offering weights were determined by observation and interviews with temple staff (Lane et al., 2011). Macaque population sizes were estimated based on multiple visits between 2000 and 2007 (Loudon et al., 2006, Lane et al., 2011).

The anthropogenic environment of Singapore was assessed previously by Klegarth (2015). Information was collected about the landscape composition of a ∼374 m radius area around various macaque sleeping locations. For sites with multiple sleeping locations, one was selected to represent the site. Population sizes were taken from Riley et al. (2015). Elevation information was collected from publicly available topographic map (Topographic Map, 2022). Sites near large water reservoirs were considered to have access to water 365 days of the year. For sites that are not near reservoirs, publicly available rainfall records from the Meteorological Service Singapore were used to determine water days (Meteorological Service Singapore, 2022). Rainfall data from the years when samples were collected was gathered and averaged. Any day with at least 6.35 mm of rain was considered a water day following Weather Insurance Agency guidelines about puddle formation (Weather Insurance Agency, 2022; Table 3).

**Table 3.** Environmental information for Singapore.

| Site | Population size | Elevation (m) | Water Days | Forest (m<sup>2</sup>) | Urban (m<sup>2</sup>) |
| --- | --- | --- | --- | --- | --- |
| AP | 26 | 16 | 146 | 186,510 | 233,938 |
| BB | 38 | 48 | 365 | 314,036 | 125,240 |
| BT | 198 | 100 | 365 | 277,394 | 123,908 |
| LP | 96 | 18 | 365 | 226,269 | 152,041 |
| MR | 325 | 30 | 365 | 442,000 | 556,740 |
| OT | 114 | 39 | 365 | 221,477 | 174,711 |
| US | 69 | 20 | 365 | 199,996 | 120,724 |
| WW | 15 | 13 | 146 | 187,922 | 63,871 |

### Linear Regressions and Model Selection

Linear regressions were performed using the “stats” package in R (v 4.2.2) to find relationships between metrics used to describe the diet of long-tailed macaques and environmental parameters associated with the sites (R Core Team, 2022). To facilitate model selection, Pearson correlations between environmental parameters were calculated. Candidate parameters were selected to avoid moderate and strong (r > 0.5) correlations between parameters. This resulted in models from Bali only considering four candidate parameters: Offering Weight (kg/day), Elevation (m), Forested Area (m^2^), and Urban Area (m^2^) (Table 4). Models from Singapore only considered three candidate parameters: Elevation (m), Urban Area (m^2^), and Water Days (Table 5).

**Table 4.** A matrix of correlations between the potential environmental parameters from Bali. Correlations with a magnitude above 0.5, considered moderate or strong correlations, are bolded. All parameters are perfectly correlated with themselves.

|  | Water Days | Tourism Days | Offering Weight | Forest Area | Rice Agriculture | Urban Area | Population Size |
| --- | --- | --- | --- | --- | --- | --- | --- |
| Tourism Days | 0.023 | 1.000 |  |  |  |  |  |
| Offering Weight | 0.241 | **0.924** | 1.000 |  |  |  |  |
| Forest Area | 0.084 | -0.450 | -0.338 | 1.000 |  |  |  |
| Rice Agriculture | **0.549** | **0.517** | **0.717** | 0.106 | 1.000 |  |  |
| Urban Area | **0.863** | -0.243 | -0.058 | 0.121 | 0.250 | 1.000 |  |
| Population Size | 0.233 | **0.665** | **0.744** | -0.434 | 0.386 | 0.065 | 1.000 |
| Elevation | -0.012 | -0.267 | -0.295 | 0.013 | -0.399 | 0.041 | -0.448 |

**Table 5.** A matrix of correlations between the potential environmental parameters from Singapore. Correlations with a magnitude above 0.5, considered moderate or strong correlations, are bolded. All parameters are perfectly correlated with themselves.

|  | Population Size | Elevation | Water Days | Forest Area |
| --- | --- | --- | --- | --- |
| Elevation | 0.391 | 1.000 |  |  |
| Water Days | **0.527** | 0.451 | 1.000 |  |
| Forest Area | **0.822** | 0.299 | 0.494 | 1.000 |
| Urban Area | **0.786** | -0.110 | 0.180 | **0.786** |

Due to the small number of sites involved in each regression, a Leave One Out Cross Validation (LOOCV) approach was employed to make sure that a single point (i.e. site) was not driving any of the regression results using the train function in the “caret” package (Kuhn, 2020). The models with the lowest Root Mean Squared Error and Mean Adjusted Error were selected and evaluated for significance. They were often, but not always, the same model, and the model with the lowest Root Mean Squared Error was selected when these metrics did not agree.

### Data Availability Statement

The data that contributed to these analyses and the R code used to run them are available as supplemental files with the publication.

## Results

### Overview of Sequencing Results

After filtering we detected 16,187,196 reads that could be assigned to a potential dietary genus and across 64 samples from Bali. We detected 14,230,152 reads that could be assigned to a potential dietary genus across 63 samples from Singapore. Overall, our reads were assigned to 1,201 potential dietary genera (Invertebrates: 665, Embryophytes: 536). Most reads were assigned to embryophyte genera (Bali: 72.32%, Singapore: 90.41%), with the remaining reads being invertebrates (Bali: 27.68%, Singapore: 9.59%). Several embryophyte genera are known macaque diet items (Supplemental Table 3; Wheatley, 1980; Corlett & Lucas, 1990; Lucas & Corlett, 1998; Sha & Honya, 2013; Brotcorne, 2014; Wheatley, 1996; Wheatley, 1999; Yeager, 1996; Ruslin et al., 2019).

### Diet Metrics for Bali

Various metrics to describe the macaque diet were calculated. It is noteworthy that sites with a single sample (AN and BT) only have one sample contributing to their average values, the average diet will equal the majority diet, and diet similarity cannot be calculated for them. The average dietary richness ranged from 33 dietary genera at Batur (BT) to 214.67 at Batu Pageh (BP). A total of 864 dietary genera were found in samples from Bali (Supplemental Table 4). The average crop dietary genera richness ranged from 8.57 at Ubud (PU) to 36.33 at Batu Pageh (BP), and 58 crop genera were detected across all samples from Bali. The average percentage of diet items that are crops ranged from 15.40% at Mekori (MK) to 27.27% at Batur (BT). The richness of the majority diet ranged from 31 dietary genera at Ubud (PU) to 186 dietary genera at Batu Pageh (BP). The similarity of the diet across samples at each site ranged from 0.10 at Uluwatu (UL) and Pulaki (PL) to 0.25 at Batu Pageh (BP; Table 6).

**Table 6.** Sample counts and diet metrics for sites in Bali, Indonesia.

| Site | Sample Count | Avg. Diet Richness | Avg. Crop Richness | Avg. Crop Percentage | Majority Diet Richness | Similarity |
| --- | --- | --- | --- | --- | --- | --- |
| AN | 1 | 51.00 | 12.00 | 23.53 | 51.00 | N/A |
| AS | 2 | 97.50 | 19.50 | 19.71 | 157.00 | 0.11 |
| BG | 8 | 109.38 | 19.38 | 18.89 | 97.00 | 0.13 |
| BP | 3 | 214.67 | 36.33 | 17.13 | 186.00 | 0.25 |
| BT | 1 | 33.00 | 9.00 | 27.27 | 33.00 | N/A |
| CK | 6 | 137.33 | 22.50 | 17.31 | 126.00 | 0.16 |
| LM | 6 | 157.83 | 23.50 | 16.17 | 152.00 | 0.18 |
| MK | 7 | 108.86 | 16.86 | 15.40 | 57.00 | 0.12 |
| PL | 4 | 86.00 | 19.50 | 23.61 | 89.00 | 0.10 |
| PU | 14 | 43.71 | 8.57 | 19.55 | 31.00 | 0.05 |
| TK | 8 | 130.13 | 25.13 | 19.95 | 101.00 | 0.15 |
| UL | 4 | 87.50 | 19.00 | 22.37 | 102.00 | 0.10 |

### Linear Models for Bali

A LOOCV model selection process suggested that the best model to explain average dietary richness of sites across Bali included two parameters: Offering Weight and Elevation (Table 7). The resulting linear model was statistically significant (p = 0.004256, Adjusted R^2^ = 0.6367; Figures 2a and S3a). Note that as provisioning increases the average dietary richness decreases, and as elevation increases the average dietary richness also decreases.

**Figure 2:**
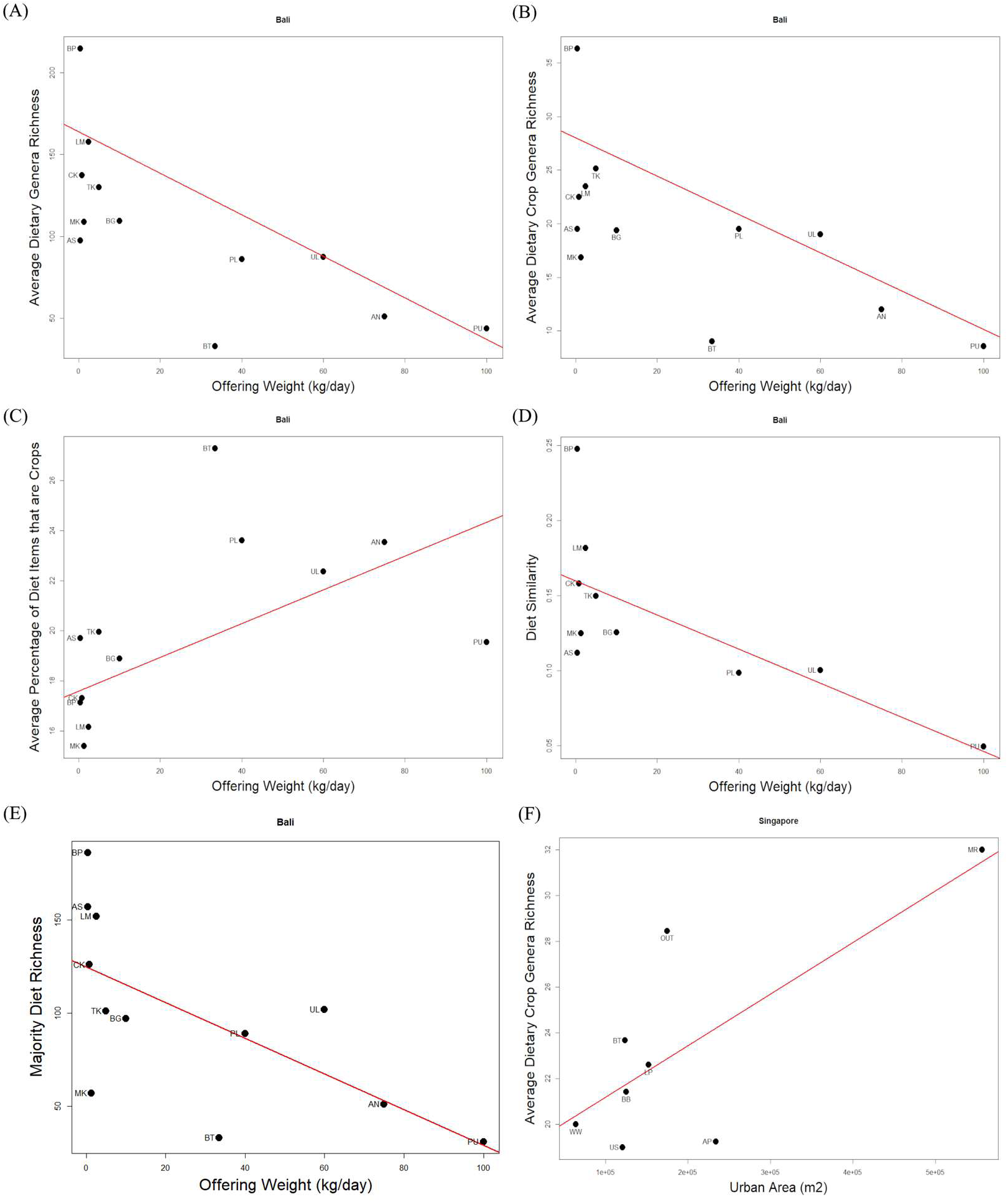
**(A)** A plot of the average dietary richness for twelve sites across Bali against the Offering Weight (kg/day). The line shows the best fit line of a statistically significant linear model (p = 0.004256; Table 7). The Adjusted R^2^ of the model is 0.6367. **(B)** A plot of the average dietary crop richness for twelve sites across Bali against the Offering Weight (kg/day). The line shows the best fit line of a statistically significant linear model (p = 0.007242; Table 7). The Adjusted R^2^ of the model is 0.5912. **(C)** A plot of the average percentage of crop contribution to dietary richness for twelve sites across Bali against the Offering Weight (kg/day). The line shows the best fit line of a statistically significant linear model (p = 0.01735; Table 7). The Adjusted R^2^ of the model is 0.5876. **(D)** A plot of the diet similarity for ten sites across Bali against the Offering Weight (kg/day). The line shows the best fit line of a statistically significant linear model (p = 0.01914; Table 7). The Adjusted R^2^ of the model is 0.4564. **(E)** A plot of the majority dietary richness for twelve sites across Bali against the Offering Weight (kg/day). The line shows the best fit line of a statistically significant linear model (p = 0.01981; Table 7). The Adjusted R^2^ of the model is 0.3775. **(F)** A plot of the average crop richness for eight sites across Singapore against the Urban Area (m^2^). The line shows the best fit line of a statistically significant linear model (p = 0.03295; Table 9). The Adjusted R^2^ of the model is 0.4855.

**Table 7.**
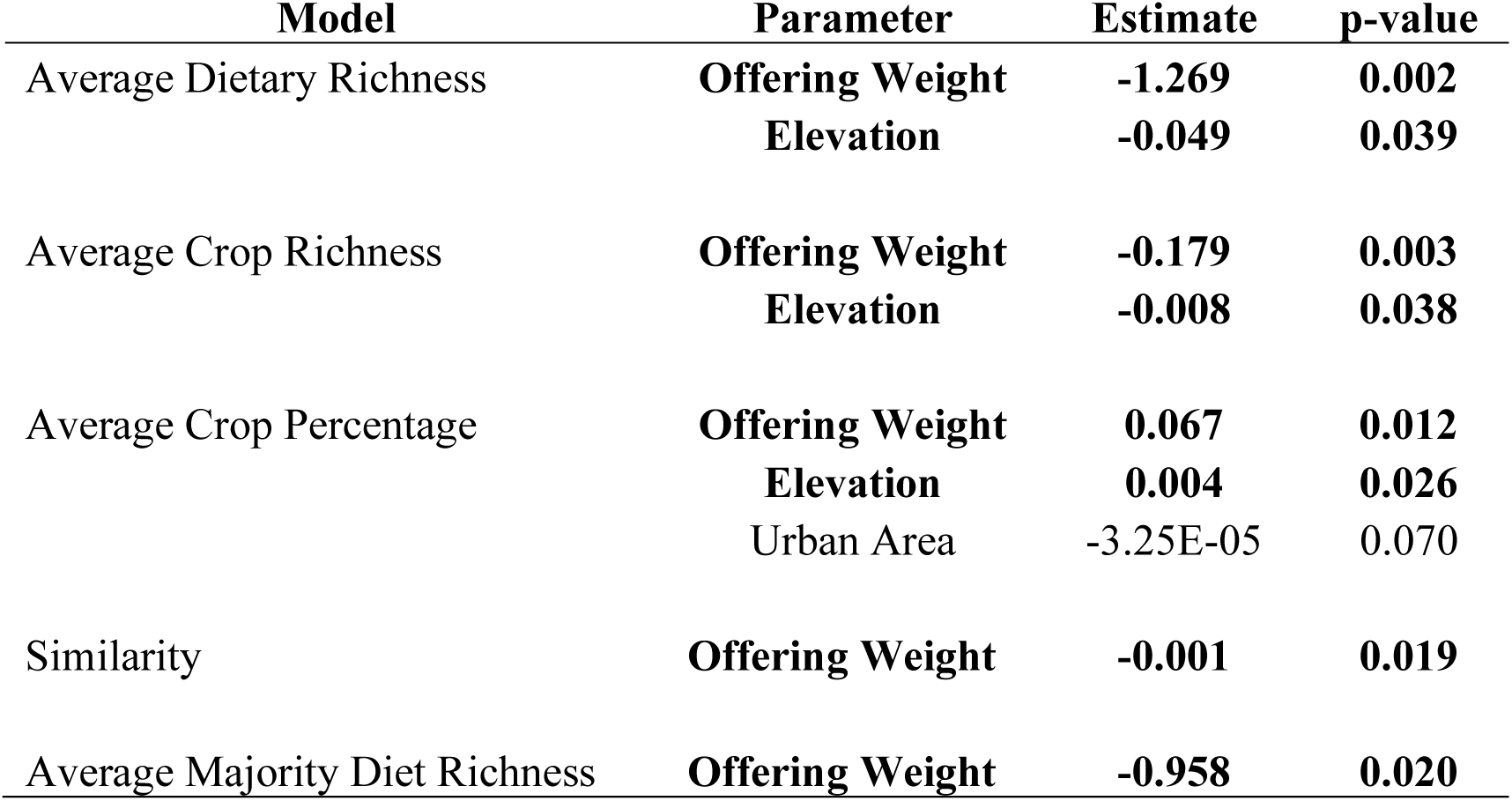
The parameters that were included in each model describing dietary metrics from Bali, including the parameter name, an estimate of the beta coefficient, and the p-value. Statistically significant parameters are bolded.

**Table 8.** Sample counts and diet metrics for sites in Singapore.

| Site | Sample Count | Avg. Diet Richness | Avg. Crop Richness | Avg. Crop Percentage | Majority Diet Richness | Similarity |
| --- | --- | --- | --- | --- | --- | --- |
| AP | 4 | 110.00 | 19.25 | 17.38 | 104.00 | 0.11 |
| BB | 12 | 125.92 | 21.42 | 19.07 | 80.00 | 0.13 |
| BT | 12 | 145.08 | 23.67 | 17.14 | 103.00 | 0.15 |
| LP | 5 | 156.80 | 22.60 | 12.70 | 118.00 | 0.16 |
| MR | 6 | 182.33 | 32.00 | 18.84 | 170.00 | 0.19 |
| OUT | 11 | 189.82 | 28.45 | 15.55 | 138.00 | 0.20 |
| US | 9 | 104.44 | 19.00 | 19.36 | 62.00 | 0.11 |
| WW | 4 | 127.25 | 20.00 | 15.73 | 149.00 | 0.13 |

**Table 9.** The parameters that were included in each model describing dietary metrics from Singapore, including the parameter name, an estimate of the beta coefficient, and the p-value. Statistically significant parameters are bolded.

| Model | Parameter | Estimate | p-value |
| --- | --- | --- | --- |
| Average Dietary Richness | Urban Area | 1.04E-04 | 0.20083 |
| Average Crop Richness | **Urban Area** | **2.06E-05** | **0.0437** |
|  | Water Days | 0.01673 | 0.2122 |
| Average Crop Percentage | Water Days | 0.002537 | 0.78802 |
| Similarity | Urban Area | 1.093E-07 | 0.200834 |
| Average Majority Diet Richness | Elevation | -0.2098 | 0.6906 |

The LOOCV model selection procedure recommended a model with two parameters to explain average dietary crop richness of sites across Bali, a measure of the number of anthropogenic plants in the diet: they were Offering Weight and Elevation (Table 7). The resulting linear model was statistically significant (p = 0.007242, Adjusted R^2^ = 0.5912; Figures 2b and S3b). As provisioning increases average dietary crop richness decreases, and as elevation increases the average dietary crop richness decreases.

The percentage of diet items that are crops, an indicator of how heavily the macaques rely on crops, was best explained by a model with Offering Weight, Elevation, and Urban Area as parameters based on the LOOCV model selection process (Table 7). The resulting linear model was statistically significant (p = 0.01735, Adjusted R^2^ = 0.5876; Figures 2c and S3c). As offering weight increases, the percentage of diet items that are crops increases and as elevation increases, the crop percentage increases.

The best model to explain diet stability (i.e. similarity) of sites across Bali only included Offering Weight as a parameter. This linear model was statistically significant (p = 0.01914, Adjusted R^2^ = 0.4564; Table 7; Figure 2d). Note that it is a negative relationship, more provisioning means that the diet for the site is less stable across individuals.

The majority diet richness for each site was best explained by a model containing a single parameter, which was Offering Weight. The LOOCV model selection process chose a statistically significant model (p = 0.01981, Adjusted R^2^ = 0.3775; Table 7; Figure 2e). As offering weight increases, majority diet richness decreases.

### Diet Metrics for Singapore

Multiple metrics were calculated to describe the diet of macaques in Singapore. Every site in Singapore is represented by at least four samples. Sites in Singapore have average dietary richness values that range from 104.44 at Upper Seletar (US) to 189.82 at Old Upper Thompson Road (OUT). A total of 951 dietary genera were found in samples from Singapore (Supplemental Table 5). For the average dietary richness of crop genera, the range is from 19.00 at Upper Seletar (US) to 32.00 at MacRitchie Nature Reserve (MR), with a total of 60 crop genera being detected in samples from Singapore. The percentage of dietary genera that are crops ranged from 12.70% at Lower Pierce (LP) to 19.36% at Upper Seletar (US). The majority dietary genera richness ranged from 62 dietary genera at Upper Seletar (US) to 170 at MacRitchie Nature Reserve (MR). The similarity of the diet at each site ranged from 0.11 at Admiralty Park (AP) and Upper Seletar (US) to 0.20 at Old Upper Thompson Road (OUT; Table 8).

### Linear Models for Singapore

The LOOCV model selection procedure was employed to identify which parameters best explained average dietary richness, diet similarity, average crop richness, average crop percentage, and majority diet genera richness. The resulting models for average diet genera richness, diet similarity, average crop percentage, and majority diet genera richness were not statistically significant (Table 9).

The remaining diet metric, average crop richness, was best explained by a model with a single parameter, which was Urban Area. The resulting model was statistically significant (p = 0.03295, Adjusted R^2^ = 0.4855; Table 9; Figure 2f). As urban area increases, crop richness increases.

## Discussion

### Bali

Long-tailed macaques are provisioned regularly at several sites across Bali. Of the environmental parameters that were tested on Bali, Offering Weight had a statistically significant impact on every dietary metric that we investigated, and it had the greatest effect in every model that included it. Offering Weight is strongly correlated with tourism days, macaque population size, and the extent of rice agriculture nearby. These correlations are easily explainable, given large amounts of provisioning facilitate large macaque populations and are associated with heavy tourism. Although we only directly tested Offering Weight, it can be considered a stand-in for the overall anthropogenic effects occurring at sites. This is in line with previous work that found that offering weight, the amount of tourism, and macaque population sizes were the parameters with the highest loading in an “anthropogenic effects” principal component (Lane et al., 2011).

The prominent role of Offering Weight in explaining every diet metric that was tested broadly suggests that anthropogenic resource access has a significant impact on the diet of long-tailed macaques in Bali.

Following the general expectation in the literature and meeting our predictions, more heavily provisioned populations have lower average dietary richness, which suggests that a relatively small number of diet items are being relied on by these macaques. The average dietary crop richness supports this conclusion. It is likely that the small subset of diet items being consumed are provisioned items, because we see the percentage of dietary genera that are crops increase as provisioning increases. Many provisioned diet items would be considered crops in our definition (Wheatley, 1980; Brotcorne, 2014; Fuentes et al., 2011). This meets our expectations that macaques with access to more anthropogenic resources will rely more heavily on crop plants. It is noteworthy that sites with very low provisioning still have a crop contribution to the diet, therefore, anthropogenic influence is ubiquitous across Bali, but highly variable in its extent and impact on diet.

Unexpectedly, as offering weight increases, the stability of the macaque diet within groups decreases. We anticipated that the temporally predictable and spatially clumped resource would result in greater stability between individuals at more heavily provisioned sites. Our opposite result may suggest that heavily provisioned sites create disparity in diet composition between individuals at a site. As a result, there may be a subset of macaques that are monopolizing access to provisioned resources, leaving other macaques to continue to forage elsewhere. Differences in “priority of access to resources” had been mentioned in previous work in Bali (Fuentes et al., 2011). Recent work in Singapore had demonstrated that sex and rank impact access to anthropogenic resources, with males and higher-ranking individuals feeding on anthropogenic food more frequently than females and lower ranking individuals, respectively (Marty, et al., 2019). The spatial clumping of resources at provisioned sites most likely facilitates this monopolization. Long-tailed macaques have a Grade 2 social system, which is characterized by unilateral aggression, highly intense aggression, and infrequent reconciliations (Thierry, 2011). This kind of social system could result in resource monopolization at sites that are heavily provisioned, explaining our seemingly contradictory results. The results regarding diet stability foreshadow unexpected results regarding majority diet richness.

In contrast to our expectations, the majority diet became less rich as offering weight increased. We had originally based this prediction on the interaction of average dietary richness decreasing and diet stability increasing with access to anthropogenic resources. Therefore, we expected the majority diet richness to increase as the macaques foraged more heavily on a small subset of provisioned foods, which would then become ubiquitous in the samples, and increase the number of diet items that were considered part of the majority diet. However, it is possible that because there was a decrease in diet stability at heavily provisioned sites due to increased competition, homogeneity of macaque diets as indicated by the number of majority diet items shared would decrease as macaques were forced to forage more broadly for whatever food items could be secured. Macaques that are monopolizing provisioned resources may have fewer diet items in common with macaques who have restricted access to those resources, resulting in fewer diet items being detected in enough samples to be considered as majority diet items.

Beyond anthropogenic impacts, Elevation was also a significant parameter influencing diet in multiple models, although to a lesser extent than Offering Weight. This relationship followed a general pattern wherein higher elevation sites had lower diet genera richness, which matches general decreases in biodiversity with increasing elevation (Dossa et al., 2013; Culmsee et al., 2010). However, this influence f rom the physical environment was always overshadowed by the influence of anthropogenic effects, highlighting their importance in anthropogenically altered environments.

### Singapore

Our model selection process did not detect any significant relationships between environmental parameters and dietary richness in samples from Singapore. This was surprising, considering that Sha & Hanya (2013) had observed an increase in diet richness with increased anthropogenic resource access. Their observations were carried out across an entire year, and whereas they saw increased diet richness with increased anthropogenic resource access across the year, this pattern was not visible when month to month comparisons were run. Our results are more in line with their month-to-month comparisons than their annual comparison (Sha & Hanya, 2013).

Although diet richness did not change, the composition of the diet did shift across Singapore as the model selection process found that urban area predicted average crop richness. This was in line with our prediction that macaques with more anthropogenic resource access will rely on a range of crop plants, and matches Sha & Hanya’s (2013) observations that macaques with more anthropogenic resource access spent more time foraging on anthropogenic foods.

Although we do not have a direct measurement of anthropogenic resource access for Singapore, urban area is likely a good proxy for anthropogenic resource access. Macaques in Singapore primarily get access to anthropogenic foods through refuse sites (e.g. garbage bins) and direct provisioning from individuals in cars (Sha & Hanya, 2013), both of which most likely occur more regularly in urban areas.

There were no statistically significant models associated with diet stability or majority diet richness in samples from Singapore. We anticipated a decrease in both stability and majority diet richness as macaques had greater access to anthropogenic resources. These predictions were partially based on Sha & Hanya’s (2013) annual observations of diet richness that were not significantly different at the monthly time scale. While the majority diet richness and stability values vary across our sites in Singapore, none of the environmental parameters that were included in our model selection process have significant impacts on them.

### How does wildlife respond to anthropogenic resources?

In general, we can conclude that anthropogenic resource access does impact the diet of long-tailed macaques. Samples from Bali suggest that increased access to anthropogenic resources results in the macaques focusing on a smaller number of dietary genera that are disproportionately crops and some individuals are likely monopolizing these anthropogenic resources. Samples from Singapore suggest that increased access to anthropogenic resources results in macaques broadening their diet to include a wider range of crops. Macaques in both island contexts increase their usage of crop diet items with greater anthropogenic access but with different resulting patterns. The lack of an overall trend is broadly in line with what Gamez et al. (2022) observed in their meta-analysis of predators living in urban environments. They found that there was not a statistically significant general trend regarding how urban environments, which involve anthropogenic resource access, interact with the dietary richness of predators.

Furthermore, they saw opposing non-significant trends in different groups of animals, with fish, birds and reptiles showing a decrease in diet richness while mammals and amphibians showed an increase in diet richness. Our results go beyond the idea that trends may vary by class of vertebrate and suggest that the response can vary even within a single species. It is likely that the contrasting patterns that we observed within long-tailed macaques are related to different properties of the anthropogenic resource. Bali’s temporally consistent and spatially clumped provisioning regime produced a different change in macaques’ diets than Singapore’s more spatially dispersed and temporally inconsistent anthropogenic resources. Gamez et al (2022) noted that there are taxonomic and geographic biases in this field, with birds, mammals, and North American species being overrepresented. While it is important to address these biases to develop stronger explanations for how wildlife species respond to anthropogenic resources, our work suggests that researchers should focus on variation in the life history details of the wildlife species (e.g. generalists, social system, etc.) *and* the anthropogenic resource (e.g. degree of spatial clumping and/or temporal predictability) as they select study systems and make predictions to further our understanding of this human/wildlife interaction.

## Supporting information

Analysis Data

Analysis Data

R Code for Analyses

R Code for Analyses

## Acknowledgments

This research was supported by the Notre Dame Center for Research Computing through use of the high-performance computing cluster. The Notre Dame Genomics and Bioinformatics Core Facility assisted through sequencing consultation and library preparation. We specifically acknowledge the contributions of Jaqueline Lopez and Melissa Stephens in the core facility for their advice and helpful suggestions regarding sequencing. We would like to acknowledge Amy Klegarth, Justin Wilcox, and Kelly Lane-deGraaf for assistance in data collection and sample processing. We would like to acknowledge Bailee Egan for assistance with sample processing. We would like to acknowledge Agustin Fuentes for his role in securing funding and samples.

Work in Bali was funded by the National Science Foundation (NSF) Division of Behavioral and Cognitive Sciences (grant BSC-0629787) and East Asian and Pacific Summer Institute Program, the Leakey Foundation, the University of Notre Dame Institute for Scholarship in the Liberal Arts, and Eck Institute for Global Health.

Work in Singapore was funded by the National Science Foundation IGERT GLOBES program (#0504495), the National Science Foundation EAPSI Singapore program 2012, the National Geographic Young Explorer’s Grant (9234-12), the US Student Fulbright Program Singapore 2013-2014, and the following groups within the University of Notre Dame: the Institute for Scholarship in the Liberal Arts, the Center for Aquatic Conservation, the College of Science, and. HSIRB (protocol numbers 07-152, 08-036, 08-039), CDC protocol numbers (2006-10-148, 2011-01-111, 2012-06-034, 2014-10-096). Samples were collected with permission from the National Parks Board (permit #NP/RP11-029) and exported under CITES (11SG006180CE, 12SG006486CE, 14SG009020CE).

## Author Contributions

BG prepared filtered tables for analyses. BG and HH planned analyses. BG executed analyses. BG and HH wrote the manuscript. CR extracted DNA, prepared samples for sequencing, and ran data through the bioinformatics pipeline. BG and AA assembled the table of environmental parameters for Singapore. All authors reviewed the manuscript.

