## Supplemental Information for "Contrasting Impacts of the Anthropogenic Environment on the Diet of Long-tailed Macaques (*Macaca fascicularis*) in Southeast Asia"

Supplementary Methods

Supplemental Table 1. A breakdown of which month and year samples from Singapore were collected in

| **Year** | **Month** | **Count** |
| --- | --- | --- |
| 2011 | March | 1 |
|  | April | 1 |
|  | May | 1 |
|  | July | 13 |
|  | August | 5 |
| 2012 | April | 3 |
|  | June | 2 |
|  | July | 9 |
| 2013 | June | 2 |
|  | July | 26 |
|  | **Total** | 63 |

OTU Dietary Genera Table Preparation

Cirtwill and Hamback (2021) outline a process for selecting a “well-chosen fixed cutoff”. This process involves correcting the read counts for tag jumping. The cutoff value is selected by looking at two plots and selecting a value near the elbow/knee of the curve. The first plot shows the number of links (i.e. non-zero cells in the table) against the cutoff value. The second plot shows the percentage of links kept at the assessed cutoff value plus one against the cutoff value. The plots were evaluated and a cutoff value of six was selected for our potential dietary genera (Figure S1).

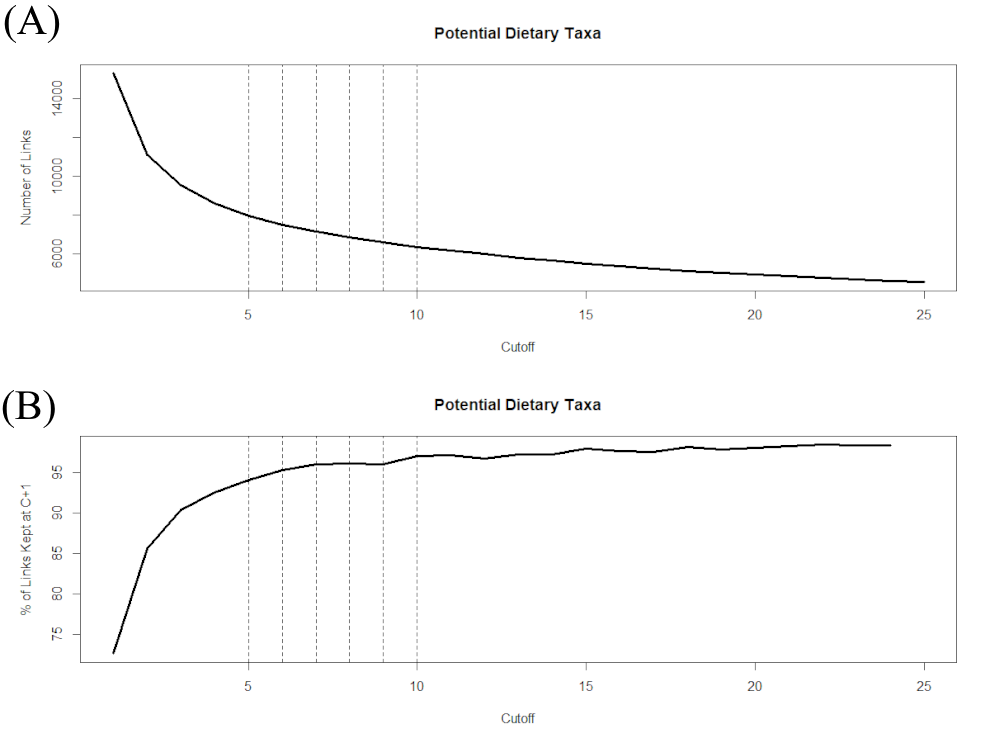

Figure S1: Example cutoff value selection plots. **(A)** A plot of the cutoff value against the number of links included in the table at that cutoff value for dietary genera detected in samples from Bali. **(B)** A plot of the cutoff value against the percentage of links kept at that cutoff value plus one for dietary genera detected in samples from Bali. Vertical dotted lines show potential cutoff values near the knee/elbow of the plots.

Majority Diet Definition

To ensure that the varying sample counts associated with each site were not predicting the majority diet richness, we used linear regressions to assess the relationship. The sample count did not predict the majority diet richness in samples from Bali (p = 0.4436) or Singapore (p = 0.2849) when a 50% threshold was used (Figure S2).

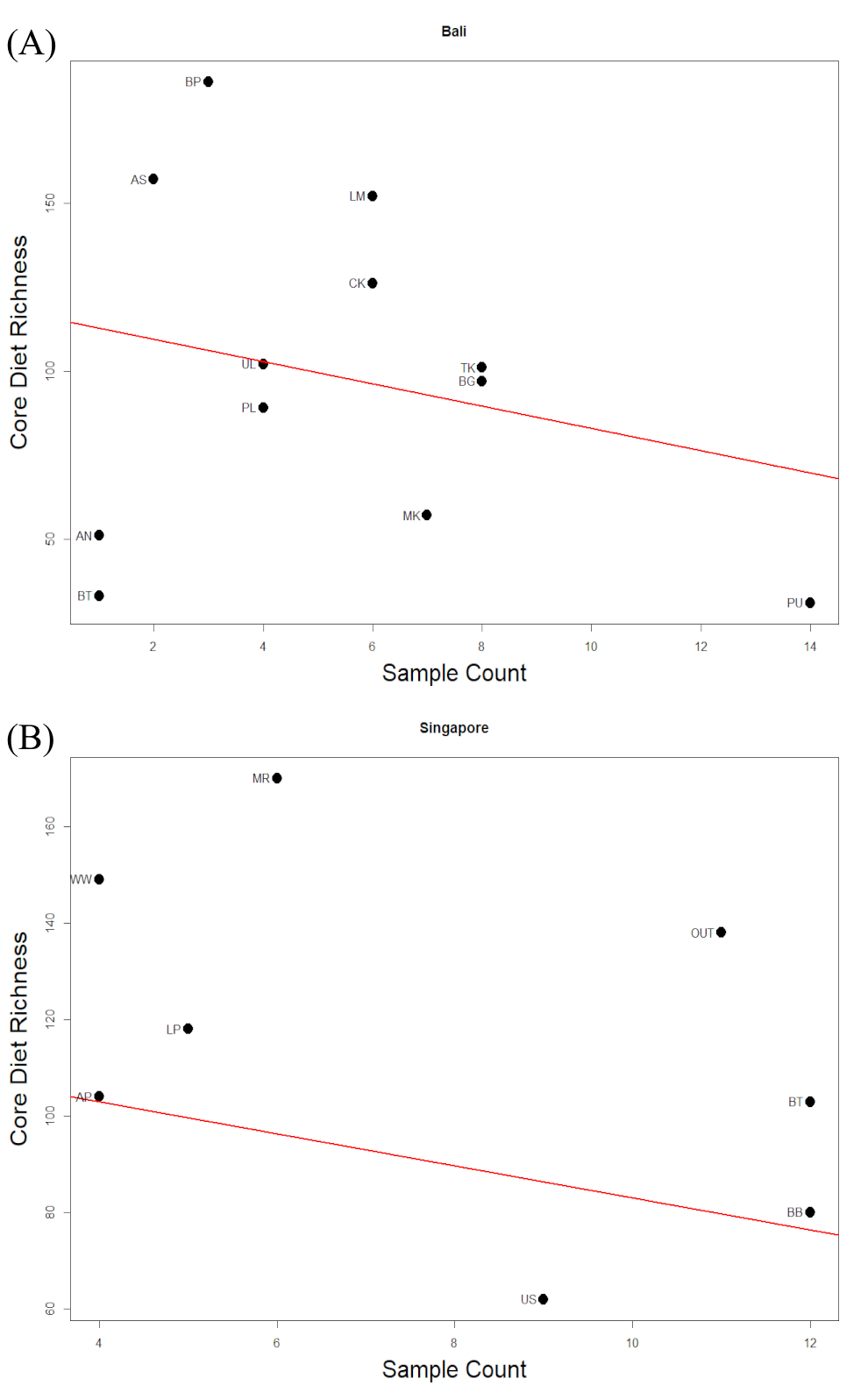

Figure S2: **(A)** A plot of the sample count against the majority diet richness for sites from Bali. The line shows the best fit line of a non-significant model (p = 0.4436). **(B)** A plot of the sample count against the majority diet richness for sites from Singapore. The line shows the best fit line of a non-significant model (p = 0.2849).

Supplemental Table 2 Note: Reprinted from Lane KE, Holley C, Hollocher H, Fuentes A. The anthropogenic environment lessens the intensity and prevalence of gastrointestinal parasites in Balinese long-tailed macaques (Macaca fascicularis). Primates. 2011 Apr;52(2):117–28 with Bedugal population omitted. Original caption: “Environmental measurements, for each macaque population, that contributed to the principal components analysis”.

| **Population** | **Water days** | **Tourism days** | **Offering weight (kg/day)** | **Forest (m^2^)** | **Rice (m^2^)** | **Urban (m^2^)** | **Population size** | **Elevation (m)** |
| --- | --- | --- | --- | --- | --- | --- | --- | --- |
| Alas Nengahn (AN) | 244 | 365 | 75 | 510781 | 724907 | 32233 | 50 | 109 |
| Angseri (AS) | 244 | 91 | 0.4 | 536319 | 123203 | 102191 | 40 | 730 |
| Bukit Gumang (BG) | 91 | 244 | 10 | 0 | 131 | 0 | 100 | 213 |
| Batu Pageh (BP) | 91 | 91 | 0.4 | 0 | 0 | 6546 | 45 | 270 |
| Batur (BT) | 91 | 244 | 33.5 | 0 | 0 | 0 | 25 | 1717 |
| Cekik (CK) | 91 | 91 | 0.8 | 977641 | 0 | 0 | 50 | 647 |
| Lempuyang (LM) | 365 | 91 | 2.5 | 180053 | 0 | 127074 | 60 | 754 |
| Mekori (MK) | 365 | 91 | 1.3 | 566899 | 298983 | 87969 | 60 | 630 |
| Pulaki (PL) | 244 | 244 | 40 | 106198 | 0 | 34488 | 200 | 679 |
| Ubud (PU) | 365 | 365 | 100 | 36002 | 807051 | 72013 | 400 | 62 |
| Tejakula (TK) | 91 | 91 | 5 | 417946 | 114393 | 23372 | 75 | 70 |
| Uluwatu (U) | 91 | 365 | 60 | 0 | 0 | 23372 | 300 | 80 |

Supplementary Results

Overview of Sequencing Results

The bioinformatics pipeline created an OTU table with 114,492,394 reads. Control samples, duplicate samples, and taxa that were not of interest were left out. OTUs in the embryophyte clade, the phylum Mollusca, or the phylum Arthropoda were put into a table of potential diet items. OTUs with enough taxonomic information were amalgamated to genera, and these genus resolution tables were split by island. These island level genus resolution tables were filtered following Cirtwill and Hamback (2021) using the previously described cutoff value of six.

Supplemental Table 3. Across nine published sources 197 dietary genera have been recorded for long-tailed macaques. We detected 43 of these previously recorded dietary genera in our data for Bali and 45 in our data for Singapore. Columns indicate where the genus was detected, with the first two columns being our metabarcoding data and the remaining ten columns being published sources. Note, Wheatley, 1996 has two columns, one for Bali (marked B) and one for Kalimantan (marked K). Numbers in the columns of published sources indicate the number of species in each dietary genus that were described in the source.

Diet Metrics for Bali

Supplemental Table 4. A total of 864 dietary genera were detected in samples from Bali, with 58 being considered crop genera. Columns indicate which genera are crop genera and which genera are considered majority dietary genera at each site. Some arthropods were labeled with the genus of the plant that they were collected from, and have been labeled as “Invertebrate” to avoid confusion (e.g. Invertebrate *Quercus*). Due to its size, this table has been included as a separate file named SupplementalTable2.xlsx.

Diet Metrics for Singapore

Supplemental Table 5. A total of 951 dietary genera were detected in samples from Singapore, with 60 being considered crop genera. Columns indicate which genera are crop genera and which genera are considered majority dietary genera at each site. Some arthropods were labeled with the genus of the plant that they were collected from, and have been labeled as “Invertebrate” to avoid confusion (e.g. Invertebrate *Quercus*). Due to its size, this table has been included as a separate file named SupplementalTable3.xlsx.

Linear Models for Bali

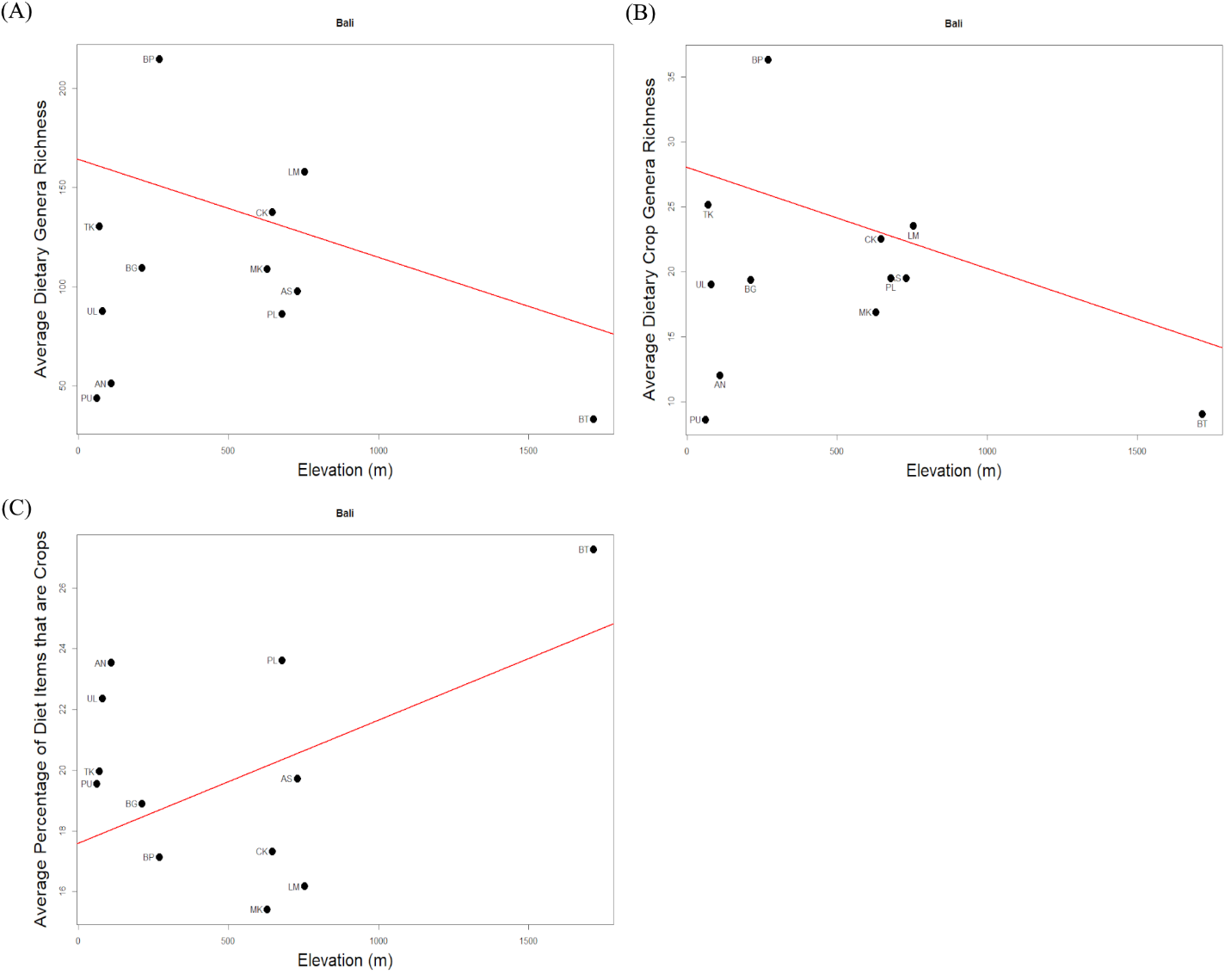

Figure S3. **(A)** A plot of the average dietary richness for twelve sites across Bali against the Elevation (m). The line shows the best fit line of a statistically significant linear model (p = 0.004256; Table 7). The Adjusted R^2^ of the model is 0.6367. **(B)** A plot of the average dietary crop richness for twelve sites across Bali against the Elevation (m). The line shows the best fit line of a statistically significant linear model (p = 0.007242; Table 7). The Adjusted R^2^ of the model is 0.5912. **(C)** A plot of the average percentage of crop contribution to dietary richness for twelve sites across Bali against the Elevation (m). The line shows the best fit line of a statistically significant linear model (p = 0.01735; Table 7).
