## Supplementary material for "Contrasting Impacts of the Anthropogenic Environment on the Diet of Long-tailed Macaques (*Macaca fascicularis*) in Southeast Asia": R Code for Analyses

#Clear the working environment
rm(list=ls())
#Load the required packages
library(caret)#This package contains functions to facilitate Leave One Out Cross Validation
library(stringr)#This package contains functions to interact with strings


#Setting the working directory, this will vary by machine
setwd("D:/01 Documents/School/ND/Macaque Diet/Publications/Diet Environment/Data")


#This is a file containing site level descriptions of the environmental parameters and diet description parameters of sites from Singapore
SingaporeData<-read.table("SingaporeData.txt",sep="\t",header=TRUE,row.names=1)

#This code calculates correlation between the environmental parameters
#Values with a magnitude between 0.5 and 0.75 are moderately correlated, while values with magnitudes above 0.75 are strongly correlated
cor(SingaporeData[,1:5])
#Based on the correlations between parameters, we are including elevation, water days, and urban area in our model selection procedure


###LOOCV Model Selection for Average Dietary Genera Richness

#This code prepares the data that will be fed into our model selection procedure
BaseFrame<-data.frame(SingaporeData[,c(6,2,3,5)])

#This code creates a series of character objects that can be combined into various model formulas
FormulaObjects<-c("AvgDietRich~", "+ Elevation", "+ Water.days", "+ Urban.Area")

#This code prepares a matrix to receive output from our model selection procedure, the Root Mean Squared Error, R Squared Value, and Mean Adjusted Error
Output<-matrix(nrow=7,ncol=3)
colnames(Output)<-c("RMSE","Rsquared","MAE")

#This code prepares a guide which sets up all possible combinations of parameters for model selection
Guide<-matrix(data=c(1,1,1,1,1,1,1,1,0,0,0,1,1,1,0,1,0,1,0,1,1,0,0,1,1,1,0,1),nrow=7,ncol=4)
colnames(Guide)<-c("AvgDietRich","Elevation","Water Days","Urban Area")
Guide

#This code sets a random seed to get consistent results
set.seed(30)

#This code runs a Leave One Out Cross Validation Model Selection procedure on every combination of parameters
#Each combination of parameters has a Root Mean Squared Error, R Squared, and Mean Adjusted Error as an output
for (i in 1:7){
  data<-BaseFrame[,which(Guide[i,]==1)]
  formula<-paste(FormulaObjects[which(Guide[i,]==1)],collapse=" ")
  formula<-str_replace(formula,pattern="\\+",replacement=" ")
  model<-train(as.formula(formula),data=BaseFrame,method="lm",
               trControl = trainControl(method = "LOOCV", number = 8, verboseIter = TRUE))
  Output[i,1:3]<-as.numeric(model$results[2:4])
}

#This code combines the output with the guide to facilitate determining which combination of parameters has the lowest RMSE
AvgDietRichOutput<-cbind(Guide[,2:4],Output)
AvgDietRichOutput

#The lowest RMSE value is 30.64636 which correlates to a model with only Urban Area as a parameter
#The lowest MAE value is 25.50161 which correlates to a model with only Urban Area as a parameter

#This code runs that linear regression
AvgDietRichModel<-lm(AvgDietRich~Urban.Area,data=SingaporeData)
summary(AvgDietRichModel)
#We can see that the model is not statistically significant


###LOOCV Model Selection for Average Dietary Crop Genera Richness

#This code prepares the data that will be fed into our model selection procedure
BaseFrame<-data.frame(SingaporeData[,c(7,2,3,5)])

#This code creates a series of character objects that can be combined into various model formulas
FormulaObjects<-c("AvgCropRich~", "+ Elevation", "+ Water.days", "+ Urban.Area")

#This code prepares a matrix to receive output from our model selection procedure, the Root Mean Squared Error, R Squared Value, and Mean Adjusted Error
Output<-matrix(nrow=7,ncol=3)
colnames(Output)<-c("RMSE","Rsquared","MAE")

#This code prepares a guide which sets up all possible combinations of parameters for model selection
Guide<-matrix(data=c(1,1,1,1,1,1,1,1,0,0,0,1,1,1,0,1,0,1,0,1,1,0,0,1,1,1,0,1),nrow=7,ncol=4)
colnames(Guide)<-c("AvgCropRich","Elevation","Water Days","Urban Area")
Guide

#This code sets a random seed to get consistent results
set.seed(30)

#This code runs a Leave One Out Cross Validation Model Selection procedure on every combination of parameters
#Each combination of parameters has a Root Mean Squared Error, R Squared, and Mean Adjusted Error as an output
for (i in 1:7){
  data<-BaseFrame[,which(Guide[i,]==1)]
  formula<-paste(FormulaObjects[which(Guide[i,]==1)],collapse=" ")
  formula<-str_replace(formula,pattern="\\+",replacement=" ")
  model<-train(as.formula(formula),data=BaseFrame,method="lm",
               trControl = trainControl(method = "LOOCV", number = 8, verboseIter = TRUE))
  Output[i,1:3]<-as.numeric(model$results[2:4])
}

#This code combines the output with the guide to facilitate determining which combination of parameters has the lowest RMSE
AvgCropRichOutput<-cbind(Guide[,2:4],Output)
AvgCropRichOutput

#The lowest RMSE value is 3.811463 which correlates to a model with Urban Area and Water Days as parameters
#The lowest MAE value is 3.076477 which correlates to a model with only Urban Area as a parameter

#This code runs the linear regression for the model that minimized RMSE
AvgCropRichModel<-lm(AvgCropRich~Urban.Area + Water.days,data=SingaporeData)
summary(AvgCropRichModel)
#We can see that the model is not statistically significant

#This code runs the linear regression for the model that minimized MAE
AvgCropRichModel<-lm(AvgCropRich~Urban.Area,data=SingaporeData)
summary(AvgCropRichModel)
#We can see that the model is statistically significant

#This code creates a plot of the significant AvgCropRichModel
par(mfrow=c(1,1),mar=c(5.1, 5.1, 4.1, 2.1))
plot(y=SingaporeData$AvgCropRich,x=SingaporeData$Urban.Area,pch = 19, ylab= "Average Dietary Crop Genera Richness",xlab="Urban Area (m2)", main = "Singapore",cex.lab =2, cex=2)
abline(AvgCropRichModel$coefficients[c(1,2)],col="red", lwd=2)
text(x = SingaporeData$Urban.Area, y = SingaporeData$AvgCropRich, labels =rownames(SingaporeData), pos=c(2,1))


###LOOCV Model Selection for Percentage of Dietary Genera which are Crops

#This code prepares the data that will be fed into our model selection procedure
BaseFrame<-data.frame(SingaporeData[,c(8,2,3,5)])

#This code creates a series of character objects that can be combined into various model formulas
FormulaObjects<-c("AvgCropPercent~", "+ Elevation", "+ Water.days", "+ Urban.Area")

#This code prepares a matrix to receive output from our model selection procedure, the Root Mean Squared Error, R Squared Value, and Mean Adjusted Error
Output<-matrix(nrow=7,ncol=3)
colnames(Output)<-c("RMSE","Rsquared","MAE")

#This code prepares a guide which sets up all possible combinations of parameters for model selection
Guide<-matrix(data=c(1,1,1,1,1,1,1,1,0,0,0,1,1,1,0,1,0,1,0,1,1,0,0,1,1,1,0,1),nrow=7,ncol=4)
colnames(Guide)<-c("AvgCropPercent","Elevation","Water Days","Urban Area")
Guide

#This code sets a random seed to get consistent results
set.seed(30)

#This code runs a Leave One Out Cross Validation Model Selection procedure on every combination of parameters
#Each combination of parameters has a Root Mean Squared Error, R Squared, and Mean Adjusted Error as an output
for (i in 1:7){
  data<-BaseFrame[,which(Guide[i,]==1)]
  formula<-paste(FormulaObjects[which(Guide[i,]==1)],collapse=" ")
  formula<-str_replace(formula,pattern="\\+",replacement=" ")
  model<-train(as.formula(formula),data=BaseFrame,method="lm",
               trControl = trainControl(method = "LOOCV", number = 8, verboseIter = TRUE))
  Output[i,1:3]<-as.numeric(model$results[2:4])
}

#This code combines the output with the guide to facilitate determining which combination of parameters has the lowest RMSE
AvgCropPercOutput<-cbind(Guide[,2:4],Output)
AvgCropPercOutput

#The lowest RMSE value is 2.600512 which correlates to a model with only Water Days as a parameter
#The lowest MAE value is 2.153295 which correlates to a model with only Urban Area as a parameter

#This code runs the linear regression for the model that minimized RMSE
AvgCropPercModel<-lm(AvgCropPercent~Water.days,data=SingaporeData)
summary(AvgCropPercModel)
#We can see that the model is not statistically significant

#This code runs the linear regression for the model that minimized MAE
AvgCropPercModel<-lm(AvgCropPercent~Urban.Area,data=SingaporeData)
summary(AvgCropPercModel)
#We can see that the model is not statistically significant

###LOOCV Model Selection for Diet Stability (i.e. Similarity)

#This code prepares the data that will be fed into our model selection procedure
BaseFrame<-data.frame(SingaporeData[,c(10,2,3,5)])

#This code creates a series of character objects that can be combined into various model formulas
FormulaObjects<-c("Similarity~", "+ Elevation", "+ Water.days", "+ Urban.Area")

#This code prepares a matrix to receive output from our model selection procedure, the Root Mean Squared Error, R Squared Value, and Mean Adjusted Error
Output<-matrix(nrow=7,ncol=3)
colnames(Output)<-c("RMSE","Rsquared","MAE")

#This code prepares a guide which sets up all possible combinations of parameters for model selection
Guide<-matrix(data=c(1,1,1,1,1,1,1,1,0,0,0,1,1,1,0,1,0,1,0,1,1,0,0,1,1,1,0,1),nrow=7,ncol=4)
colnames(Guide)<-c("Similarity","Elevation","Water Days","Urban Area")
Guide

#This code sets a random seed to get consistent results
set.seed(30)

#This code runs a Leave One Out Cross Validation Model Selection procedure on every combination of parameters
#Each combination of parameters has a Root Mean Squared Error, R Squared, and Mean Adjusted Error as an output
for (i in 1:7){
  data<-BaseFrame[,which(Guide[i,]==1)]
  formula<-paste(FormulaObjects[which(Guide[i,]==1)],collapse=" ")
  formula<-str_replace(formula,pattern="\\+",replacement=" ")
  model<-train(as.formula(formula),data=BaseFrame,method="lm",
               trControl = trainControl(method = "LOOCV", number = 8, verboseIter = TRUE))
  Output[i,1:3]<-as.numeric(model$results[2:4])
}

#This code combines the output with the guide to facilitate determining which combination of parameters has the lowest RMSE
SimilarityOutput<-cbind(Guide[,2:4],Output)
SimilarityOutput

#The lowest RMSE value is 0.03225933 which correlates to a model with only Urban Area as a parameter
#The lowest MAE value is 0.268438 which correlates to a model with only Urban Area as a parameter

#This code runs that linear regression
SimilarityModel<-lm(Similarity~Urban.Area,data=SingaporeData)
summary(SimilarityModel)
#We can see that the model is not statistically significant


###LOOCV Model Selection for Richness of the Core Diet

#This code prepares the data that will be fed into our model selection procedure
BaseFrame<-data.frame(SingaporeData[,c(9,2,3,5)])

#This code creates a series of character objects that can be combined into various model formulas
FormulaObjects<-c("CoreDietRich~", "+ Elevation", "+ Water.days", "+ Urban.Area")

#This code prepares a matrix to receive output from our model selection procedure, the Root Mean Squared Error, R Squared Value, and Mean Adjusted Error
Output<-matrix(nrow=7,ncol=3)
colnames(Output)<-c("RMSE","Rsquared","MAE")

#This code prepares a guide which sets up all possible combinations of parameters for model selection
Guide<-matrix(data=c(1,1,1,1,1,1,1,1,0,0,0,1,1,1,0,1,0,1,0,1,1,0,0,1,1,1,0,1),nrow=7,ncol=4)
colnames(Guide)<-c("CoreDietRich","Elevation","Water Days","Urban Area")
Guide

#This code sets a random seed to get consistent results
set.seed(30)

#This code runs a Leave One Out Cross Validation Model Selection procedure on every combination of parameters
#Each combination of parameters has a Root Mean Squared Error, R Squared, and Mean Adjusted Error as an output
for (i in 1:7){
  data<-BaseFrame[,which(Guide[i,]==1)]
  formula<-paste(FormulaObjects[which(Guide[i,]==1)],collapse=" ")
  formula<-str_replace(formula,pattern="\\+",replacement=" ")
  model<-train(as.formula(formula),data=BaseFrame,method="lm",
               trControl = trainControl(method = "LOOCV", number = 8, verboseIter = TRUE))
  Output[i,1:3]<-as.numeric(model$results[2:4])
}

#This code combines the output with the guide to facilitate determining which combination of parameters has the lowest RMSE
CoreDietRichOutput<-cbind(Guide[,2:4],Output)
CoreDietRichOutput

#The lowest RMSE value is 39.3077 which correlates to a model with only Elevation as a parameter
#The lowest MAE value is 32.32633 which correlates to a model with only Elevation as a parameter

#This code runs that linear regression
CoreDietRichModel<-lm(CoreDietRich~Elevation,data=SingaporeData)
summary(CoreDietRichModel)
#We can see that the model is not statistically significant
