## Supplementary material for "Contrasting Impacts of the Anthropogenic Environment on the Diet of Long-tailed Macaques (*Macaca fascicularis*) in Southeast Asia": R Code for Analyses

#Clear the working environment
rm(list=ls())
#Load the required packages
library(caret)#This package contains functions to facilitate Leave One Out Cross Validation
library(stringr)#This package contains functions to interact with strings


#Setting the working directory, this will vary by machine
setwd("D:/01 Documents/School/ND/Macaque Diet/Publications/Diet Environment/Data")


#This is a file containing site level descriptions of the environmental parameters and diet description parameters of sites from Bali
BaliData<-read.table("BaliData.txt",sep="\t",header=TRUE,row.names=1)

#This code calculates correlation between the environmental parameters
#Values with a magnitude between 0.5 and 0.75 are moderately correlated, while values with magnitudes above 0.75 are strongly correlated
cor(BaliData[,1:8])
#Based on the correlations between parameters, we are including elevation, forested area, offering weight, and urban area


###LOOCV Model Selection for Average Dietary Genera Richness

#This code prepares the data that will be fed into our model selection procedure
BaseFrame<-data.frame(BaliData[,c(9,3,4,6,8)])

#This code creates a series of character objects that can be combined into various model formulas
FormulaObjects<-c("AvgDietRich~", "+ Offering_weight_kg.day", "+ Forest_m2", "+ Urban_m2", "+ Elevation_m")

#This code prepares a matrix to receive output from our model selection procedure, the Root Mean Squared Error, R Squared Value, and Mean Adjusted Error
Output<-matrix(nrow=15,ncol=3)
colnames(Output)<-c("RMSE","Rsquared","MAE")

#This code prepares a guide which sets up all possible combinations of parameters for model selection
Guide<-matrix(data=c(1,1,1,1,1,1,1,1,1,1,1,1,1,1,1,1,0,0,0,1,1,1,0,0,0,1,1,1,0,1,0,1,0,0,1,0,0,1,1,0,1,1,0,1,1,0,0,1,0,0,1,0,1,0,1,1,0,1,1,1,0,0,0,1,0,0,1,0,1,1,0,1,1,1,1),nrow=15,ncol=5)
colnames(Guide)<-c("AvgDietRich","OfferingWeight","ForestArea","UrbanArea","Elevation")
Guide


#This code sets a random seed to get consistent results
set.seed(30)

#This code runs a Leave One Out Cross Validation Model Selection procedure on every combination of parameters
#Each combination of parameters has a Root Mean Squared Error, R Squared, and Mean Adjusted Error as an output
for (i in 1:15){
  data<-BaseFrame[,which(Guide[i,]==1)]
  formula<-paste(FormulaObjects[which(Guide[i,]==1)],collapse=" ")
  formula<-str_replace(formula,pattern="\\+",replacement=" ")
  model<-train(as.formula(formula),data=BaseFrame,method="lm",
               trControl = trainControl(method = "LOOCV", number = 12, verboseIter = TRUE))
  Output[i,1:3]<-as.numeric(model$results[2:4])
}

#This code combines the output with the guide to facilitate determining which combination of parameters has the lowest RMSE
AvgDietRichOutput<-cbind(Guide[,2:5],Output)
AvgDietRichOutput

#The lowest RMSE value is 33.23686 which correlates to a model with Offering Weight and Elevation as parameters.
#The lowest MAE value is 26.71537 which correlates to a model with Offering Weight and Elevation as parameters.

#This code runs that linear regression
AvgDietRichModel<-lm(AvgDietRich~Offering_weight_kg.day + Elevation_m,data=BaliData)
summary(AvgDietRichModel)
#We can see that the model is statistically significant

#This code creates a plot of the AvgDietRichModel with Offering Weight on the X axis
par(mfrow=c(1,1),mar=c(5.1, 5.1, 4.1, 2.1))
plot(y=BaliData$AvgDietRich,x=BaliData$Offering_weight_kg.day,pch = 19, ylab= "Average Dietary Genera Richness",xlab="Offering Weight (kg/day)", main = "Bali",cex.lab =2, cex=2)
abline(AvgDietRichModel$coefficients[c(1,2)],col="red", lwd=2)
text(x = BaliData$Offering_weight_kg.day, y = BaliData$AvgDietRich, labels =rownames(BaliData), pos=2)

#This code creates a plot of the AvgDietRichModel with Elevation on the X axis
par(mfrow=c(1,1),mar=c(5.1, 5.1, 4.1, 2.1))
plot(y=BaliData$AvgDietRich,x=BaliData$Elevation_m,pch = 19, ylab= "Average Dietary Genera Richness",xlab="Elevation (m)", main = "Bali",cex.lab =2, cex=2)
abline(AvgDietRichModel$coefficients[c(1,3)],col="red", lwd=2)
text(x = BaliData$Elevation_m, y = BaliData$AvgDietRich, labels =rownames(BaliData), pos=2)


###LOOCV Model Selection for Average Dietary Crop Genera Richness

#This code prepares the data that will be fed into our model selection procedure
BaseFrame<-data.frame(BaliData[,c(10,3,4,6,8)])

#This code creates a series of character objects that can be combined into various model formulas
FormulaObjects<-c("AvgCropRich~", "+ Offering_weight_kg.day", "+ Forest_m2", "+ Urban_m2", "+ Elevation_m")

#This code prepares a matrix to receive output from our model selection procedure, the Root Mean Squared Error, R Squared Value, and Mean Adjusted Error
Output<-matrix(nrow=15,ncol=3)
colnames(Output)<-c("RMSE","Rsquared","MAE")

#This code prepares a guide which sets up all possible combinations of parameters for model selection
Guide<-matrix(data=c(1,1,1,1,1,1,1,1,1,1,1,1,1,1,1,1,0,0,0,1,1,1,0,0,0,1,1,1,0,1,0,1,0,0,1,0,0,1,1,0,1,1,0,1,1,0,0,1,0,0,1,0,1,0,1,1,0,1,1,1,0,0,0,1,0,0,1,0,1,1,0,1,1,1,1),nrow=15,ncol=5)
colnames(Guide)<-c("AvgDietRich","OfferingWeight","ForestArea","UrbanArea","Elevation")
Guide


#This code sets a random seed to get consistent results
set.seed(30)

#This code runs a Leave One Out Cross Validation Model Selection procedure on every combination of parameters
#Each combination of parameters has a Root Mean Squared Error, R Squared, and Mean Adjusted Error as an output
for (i in 1:15){
  data<-BaseFrame[,which(Guide[i,]==1)]
  formula<-paste(FormulaObjects[which(Guide[i,]==1)],collapse=" ")
  formula<-str_replace(formula,pattern="\\+",replacement=" ")
  model<-train(as.formula(formula),data=BaseFrame,method="lm",
               trControl = trainControl(method = "LOOCV", number = 12, verboseIter = TRUE))
  Output[i,1:3]<-as.numeric(model$results[2:4])
}

#This code combines the output with the guide to facilitate determining which combination of parameters has the lowest RMSE
AvgCropRichOutput<-cbind(Guide[,2:5],Output)
AvgCropRichOutput

#The lowest RMSE value is 5.111884 which correlates to a model with Offering Weight and Elevation as parameters
#The lowest MAE value is 3.926406 which correlates to a model with Offering Weight and Elevation as parameters

#This code runs that linear regression
AvgCropRichModel<-lm(AvgCropRich~Offering_weight_kg.day + Elevation_m,data=BaliData)
summary(AvgCropRichModel)
#We can see that the model is statistically significant

#This code creates a plot of the AvgCropRichModel
par(mfrow=c(1,1),mar=c(5.1, 5.1, 4.1, 2.1))
plot(y=BaliData$AvgCropRich,x=BaliData$Offering_weight_kg.day,pch = 19, ylab= "Average Dietary Crop Genera Richness",xlab="Offering Weight (kg/day)", main = "Bali",cex.lab =2, cex=2)
abline(AvgCropRichModel$coefficients[c(1,2)],col="red", lwd=2)
text(x = BaliData$Offering_weight_kg.day, y = BaliData$AvgCropRich, labels =rownames(BaliData), pos=c(1,2))

#This code creates a plot of the AvgCropRichModel with Elevation on the X axis
par(mfrow=c(1,1),mar=c(5.1, 5.1, 4.1, 2.1))
plot(y=BaliData$AvgCropRich,x=BaliData$Elevation_m,pch = 19, ylab= "Average Dietary Crop Genera Richness",xlab="Elevation (m)", main = "Bali",cex.lab =2, cex=2)
abline(AvgCropRichModel$coefficients[c(1,3)],col="red", lwd=2)
text(x = BaliData$Elevation_m, y = BaliData$AvgCropRich, labels =rownames(BaliData), pos= c(1,2))


###LOOCV Model Selection for Percentage of Dietary Genera which are Crops

#This code prepares the data that will be fed into our model selection procedure
BaseFrame<-data.frame(BaliData[,c(11,3,4,6,8)])

#This code creates a series of character objects that can be combined into various model formulas
FormulaObjects<-c("AvgCropPercent~", "+ Offering_weight_kg.day", "+ Forest_m2", "+ Urban_m2", "+ Elevation_m")

#This code prepares a matrix to receive output from our model selection procedure, the Root Mean Squared Error, R Squared Value, and Mean Adjusted Error
Output<-matrix(nrow=15,ncol=3)
colnames(Output)<-c("RMSE","Rsquared","MAE")

#This code prepares a guide which sets up all possible combinations of parameters for model selection
Guide<-matrix(data=c(1,1,1,1,1,1,1,1,1,1,1,1,1,1,1,1,0,0,0,1,1,1,0,0,0,1,1,1,0,1,0,1,0,0,1,0,0,1,1,0,1,1,0,1,1,0,0,1,0,0,1,0,1,0,1,1,0,1,1,1,0,0,0,1,0,0,1,0,1,1,0,1,1,1,1),nrow=15,ncol=5)
colnames(Guide)<-c("AvgDietRich","OfferingWeight","ForestArea","UrbanArea","Elevation")
Guide


#This code sets a random seed to get consistent results
set.seed(30)

#This code runs a Leave One Out Cross Validation Model Selection procedure on every combination of parameters
#Each combination of parameters has a Root Mean Squared Error, R Squared, and Mean Adjusted Error as an output
for (i in 1:15){
  data<-BaseFrame[,which(Guide[i,]==1)]
  formula<-paste(FormulaObjects[which(Guide[i,]==1)],collapse=" ")
  formula<-str_replace(formula,pattern="\\+",replacement=" ")
  model<-train(as.formula(formula),data=BaseFrame,method="lm",
               trControl = trainControl(method = "LOOCV", number = 12, verboseIter = TRUE))
  Output[i,1:3]<-as.numeric(model$results[2:4])
}


#This code combines the output with the guide to facilitate determining which combination of parameters has the lowest RMSE
AvgCropPercOutput<-cbind(Guide[,2:5],Output)
AvgCropPercOutput

#The lowest RMSE value is 2.969660 which correlates to a model with Offering Weight, Urban Area, and Elevation as parameters
#The lowest MAE value is 2.588003 which correlates to a model with Offering Weight, Urban Area, and Elevation as parameters

#This code runs that linear regression
AvgCropPercModel<-lm(AvgCropPercent~Offering_weight_kg.day + Elevation_m + Urban_m2,data=BaliData)
summary(AvgCropPercModel)
#We can see that the model is statistically significant with urban area as a non-significant parameter

#This code creates a plot of the AvgCropPercModel
par(mfrow=c(1,1),mar=c(5.1, 5.1, 4.1, 2.1))
plot(y=BaliData$AvgCropPerc,x=BaliData$Offering_weight_kg.day,pch = 19, ylab= "Average Percentage of Diet Items that are Crops",xlab="Offering Weight (kg/day)", main = "Bali",cex.lab =2, cex=2)
abline(AvgCropPercModel$coefficients[c(1,2)],col="red", lwd=2)
text(x = BaliData$Offering_weight_kg.day, y = BaliData$AvgCropPerc, labels =rownames(BaliData), pos=2)

#This code creates a plot of the AvgCropPercModel with Elevation on the X axis
par(mfrow=c(1,1),mar=c(5.1, 5.1, 4.1, 2.1))
plot(y=BaliData$AvgCropPerc,x=BaliData$Elevation_m,pch = 19, ylab= "Average Percentage of Diet Items that are Crops",xlab="Elevation (m)", main = "Bali",cex.lab =2, cex=2)
abline(AvgCropPercModel$coefficients[c(1,3)],col="red", lwd=2)
text(x = BaliData$Elevation_m, y = BaliData$AvgCropPerc, labels =rownames(BaliData), pos=2)


###LOOCV Model Selection for Diet Similarity (i.e. Stability)

#This code prepares the data that will be fed into our model selection procedure
#Note that sites AN and BT have only one sample, which means that there is no way to calculate similarity at the site
#These sites were dropped from this analysis
BaseFrame<-data.frame(BaliData[-c(1,5),c(13,3,4,6,8)])

#This code creates a series of character objects that can be combined into various model formulas
FormulaObjects<-c("Similarity~", "+ Offering_weight_kg.day", "+ Forest_m2", "+ Urban_m2", "+ Elevation_m")

#This code prepares a matrix to receive output from our model selection procedure, the Root Mean Squared Error, R Squared Value, and Mean Adjusted Error
Output<-matrix(nrow=15,ncol=3)
colnames(Output)<-c("RMSE","Rsquared","MAE")

#This code prepares a guide which sets up all possible combinations of parameters for model selection
Guide<-matrix(data=c(1,1,1,1,1,1,1,1,1,1,1,1,1,1,1,1,0,0,0,1,1,1,0,0,0,1,1,1,0,1,0,1,0,0,1,0,0,1,1,0,1,1,0,1,1,0,0,1,0,0,1,0,1,0,1,1,0,1,1,1,0,0,0,1,0,0,1,0,1,1,0,1,1,1,1),nrow=15,ncol=5)
colnames(Guide)<-c("Similarity","+ Offering_weight_kg.day", "+ Forest_m2", "+ Urban_m2", "+ Elevation_m")
Guide

#This code sets a random seed to get consistent results
set.seed(30)

#This code runs a Leave One Out Cross Validation Model Selection procedure on every combination of parameters
#Each combination of parameters has a Root Mean Squared Error, R Squared, and Mean Adjusted Error as an output
for (i in 1:15){
  data<-BaseFrame[,which(Guide[i,]==1)]
  formula<-paste(FormulaObjects[which(Guide[i,]==1)],collapse=" ")
  formula<-str_replace(formula,pattern="\\+",replacement=" ")
  model<-train(as.formula(formula),data=BaseFrame,method="lm",
               trControl = trainControl(method = "LOOCV", number = 10, verboseIter = TRUE))
  Output[i,1:3]<-as.numeric(model$results[2:4])
}

#This code combines the output with the guide to facilitate determining which combination of parameters has the lowest RMSE
SimilarityOutput<-cbind(Guide[,2:5],Output)
SimilarityOutput

#The lowest RMSE value is 0.04162130 which correlates to a model with only Offering Weight as a parameter
#The lowest MAE value is 0.02982137 which correlates to a model with only Offering Weight as a parameter

#This code runs that linear regression
SimilarityModel<-lm(Similarity~Offering_weight_kg.day,data=BaliData)
summary(SimilarityModel)
#We can see that the model is statistically significant

#This code creates a plot of the SimilarityModel
par(mfrow=c(1,1),mar=c(5.1, 5.1, 4.1, 2.1))
plot(y=BaliData$Similarity,x=BaliData$Offering_weight_kg.day,pch = 19, ylab= "Diet Similarity",xlab="Offering Weight (kg/day)", main = "Bali",cex.lab =2, cex=2)
abline(SimilarityModel$coefficients[c(1,2)],col="red", lwd=2)
text(x = BaliData$Offering_weight_kg.day, y = BaliData$Similarity, labels =rownames(BaliData), pos=2)


###LOOCV Model Selection for Richness of the Majority Diet

#This code prepares the data that will be fed into our model selection procedure
BaseFrame<-data.frame(BaliData[,c(12,3,4,6,8)])

#This code creates a series of character objects that can be combined into various model formulas
FormulaObjects<-c("CoreDietRich~", "+ Offering_weight_kg.day", "+ Forest_m2", "+ Urban_m2", "+ Elevation_m")

#This code prepares a matrix to receive output from our model selection procedure, the Root Mean Squared Error, R Squared Value, and Mean Adjusted Error
Output<-matrix(nrow=15,ncol=3)
colnames(Output)<-c("RMSE","Rsquared","MAE")

#This code prepares a guide which sets up all possible combinations of parameters for model selection
Guide<-matrix(data=c(1,1,1,1,1,1,1,1,1,1,1,1,1,1,1,1,0,0,0,1,1,1,0,0,0,1,1,1,0,1,0,1,0,0,1,0,0,1,1,0,1,1,0,1,1,0,0,1,0,0,1,0,1,0,1,1,0,1,1,1,0,0,0,1,0,0,1,0,1,1,0,1,1,1,1),nrow=15,ncol=5)
colnames(Guide)<-c("CoreDietRich","OfferingWeight","ForestArea","UrbanArea","Elevation")
Guide


#This code sets a random seed to get consistent results
set.seed(30)

#This code runs a Leave One Out Cross Validation Model Selection procedure on every combination of parameters
#Each combination of parameters has a Root Mean Squared Error, R Squared, and Mean Adjusted Error as an output
for (i in 1:15){
  data<-BaseFrame[,which(Guide[i,]==1)]
  formula<-paste(FormulaObjects[which(Guide[i,]==1)],collapse=" ")
  formula<-str_replace(formula,pattern="\\+",replacement=" ")
  model<-train(as.formula(formula),data=BaseFrame,method="lm",
               trControl = trainControl(method = "LOOCV", number = 12, verboseIter = TRUE))
  Output[i,1:3]<-as.numeric(model$results[2:4])
}


#This code combines the output with the guide to facilitate determining which combination of parameters has the lowest RMSE
CoreDietRichOutput<-cbind(Guide[,2:4],Output)
CoreDietRichOutput

#The lowest RMSE value is 41.38243 which correlates to a model with only Offering Weight as a parameter
#The lowest MAE value is 31.75302 which correlates to a model with only Offering Weight as a parameter

#This code runs that linear regression
CoreDietRichModel<-lm(CoreDietRich~Offering_weight_kg.day,data=BaliData)
summary(CoreDietRichModel)
#We can see that the model is statistically significant

#This code creates a plot of the CoreDietRichModel
par(mfrow=c(1,1),mar=c(5.1, 5.1, 4.1, 2.1))
plot(y=BaliData$CoreDietRich,x=BaliData$Offering_weight_kg.day,pch = 19, ylab= "Majority Diet Richness",xlab="Offering Weight (kg/day)", main = "Bali",cex.lab =2, cex=2)
abline(CoreDietRichModel$coefficients[c(1,2)],col="red", lwd=2)
text(x = BaliData$Offering_weight_kg.day, y = BaliData$CoreDietRich, labels =rownames(BaliData), pos=2)
